# Hypnotic visual hallucination induces greater lateralised brain activity than visual imagery

**DOI:** 10.1101/2021.03.04.434014

**Authors:** Renzo C. Lanfranco, Álvaro Rivera-Rei, David Huepe, Agustín Ibáñez, Andrés Canales-Johnson

**Affiliations:** Department of Neuroscience, Karolinska Institute, Stockholm, Sweden; Department of Psychology, University of Edinburgh, Edinburgh, United Kingdom; Latin American Brain Health Institute (BrainLat) & Center for Social and Cognitive Neuroscience, Universidad Adolfo Ibáñez, Santiago, Chile; Cognitive Neuroscience Center, Universidad de San Andrés, Buenos Aires, Argentina; National Scientific and Technical Research Council (CONICET), Buenos Aires, Argentina; Global Brain Health Institute, University of California San Francisco, San Francisco, United States of America, and Trinity College Dublin, Dublin, Ireland; Department of Psychology, University of Cambridge, Cambridge, United Kingdom; Vicerrectoría de Investigación y Posgrado, Universidad Católica del Maule, Talca, Chile

## Abstract

Hypnotic suggestions can produce a broad range of perceptual experiences, including hallucinations. Visual hypnotic hallucinations differ in many ways from regular mental images. For example, they are usually experienced as automatic, vivid, and real images, typically compromising the sense of reality. While both hypnotic hallucination and mental imagery are believed to mainly rely on the activation of the visual cortex via top-down mechanisms, it is unknown how they differ in the neural processes they engage. Here we used an adaptation paradigm to test and compare top-down processing between hypnotic hallucination, mental imagery, and visual perception in very highly hypnotisable individuals whose ability to hallucinate was assessed. By measuring the N170/VPP event-related complex and using multivariate decoding analysis, we found that hypnotic hallucination of faces involves greater top-down activation of sensory processing through lateralised mechanisms in the right hemisphere compared to mental imagery. Our findings suggest that the neural signatures that distinguish hypnotically hallucinated faces from imagined faces lie in the right brain hemisphere.

## 1 Introduction

Hypnotic suggestions are suggested changes in perception, cognition, or behaviour typically preceded by a hypnotic induction procedure (Oakley & Halligan, 2013). The ability to experience hallucinations in response to hypnotic suggestions is one of the hallmarks of high hypnotisable people (Canales-Johnson et al., 2012; Hilgard, 1965; Lynn et al., 2010; Nash & Barnier, 2012). Hypnotic hallucinations are described as effortless, spontaneous, vivid, and automatic changes in perceptual experience. Oftentimes, they compromise the sense of reality. Multiple studies have explored the neural underpinnings of hypnotic hallucination in vision (Kosslyn et al., 2000; Mazzoni et al., 2009; McGeown et al., 2012; Schmidt et al., 2017; Spiegel et al., 1985), audition (Franz et al., 2020; Szechtman et al., 1998), and somatosensory processing (Derbyshire et al., 2004; Perri et al., 2019). However, the nature of hypnotic hallucinations remains poorly understood. Critically, it is unclear what neural processes distinguish hypnotic hallucination from mental imagery.

Hypnotic hallucination differs in many ways from imagination (Fazekas, 2021; Waters et al., 2021). While hypnotic hallucinations are experienced as automatic, involuntary, spontaneous, and real (Bowers, 1967; Bowers & Gilmore, 1969; Hilgard, 1965), mental images are effortful, goal-directed, and easily discernible as fictional (Canales-Johnson et al., 2021; Kosslyn, 2005; Thompson, 2007). Since neither hypnotic hallucination nor mental imagery seem to require sensory inputs to occur, they are believed to differ in their top-down influence on sensory cortices (Oakley & Halligan, 2013; Pearson, 2019; Powers et al., 2016; for a review, see Terhune et al., 2017). However, if this is the case, then how do these two processes differ from each other, and regular perception, in how they create a perceptual experience?

Hypnotic hallucinations involve changes in brain function similar to those that occur in perception. For example, Kosslyn et al. (2000) reported that highly hypnotisable participants are able to perceive a grey pattern as coloured (and vice versa) when instructed to do so during hypnosis. Using Positron Emission Tomography, they found that hypnotic hallucinations of colour involve bilateral changes in activation of the fusiform gyrus. These findings have been replicated on their subjective ratings of colour (Mazzoni et al., 2009) and brain activity changes (McGeown et al., 2012), in particular in studies with hypnotic virtuosos (i.e. very responsive highly hypnotisable individuals; Kallio & Koivisto, 2013, 2016; Koivisto et al., 2013). Other studies have investigated the ability of hypnotic hallucinations to obstruct perceptual processing. For example, Spiegel et al. (1985) reported that high hypnotisable participants show significantly lower amplitude in the N2 and P3 event-related potential (ERP) components (which are associated with selective attention) when instructed to see a cardboard box blocking the view of the monitor screen (also see Barabasz & Lonsdale, 1983; Jasiukaitis et al., 1996). More recently, a similar study found that hypnotic hallucinations of blockage not only alter ERP components but also impair counting performance in a visual identification task (Schmidt et al., 2017). Thus, hypnotic hallucinations can modulate perceptual processing following suggestions.

Similar to hypnotic hallucination, mental imagery is also believed to activate similar neural processes and representations than perception (Borst & Kosslyn, 2008; Ganis et al., 2004; Ishai et al., 2000; Kosslyn, 1996; Kosslyn et al., 2006), however, mental imagery and perception differ in how they engage with bottom-up and top-down mechanisms (Dijkstra et al., 2017; Ganis & Schendan, 2008; also see Koenig-Robert & Pearson, 2021). For example, Ganis & Schendan (2008) measured the ERP complex N170/VPP, a face-sensitive visual evoked potential (Bentin et al., 1996; Eimer, 2011), in response to face images (test stimuli) that were preceded either by a perceived or an imagined face image (adaptor stimuli). They found that while perceived adaptors suppressed the ERP’s amplitude to test stimuli, imagined adaptors enhanced it. Since mental imagery, unlike perception, entails a reactivation of visual representations through memory recall and attention, these results were interpreted as evidence of top-down activation by mental imagery. This adaptation approach has been repeatedly used to study neural representations (e.g. Amihai et al., 2011; Kaiser et al., 2013; for a review: Grill-Spector et al., 2006). The logic behind it is that repeated stimuli yield suppression effects because: (a) neurons that respond to the stimulus presentation enter a fatigue stage thus decreasing their firing rate; (b) firing neurons become sparser as a consequence of a weaker response from neurons that code less relevant features; and (c) firing neurons become more efficient at responding to the same stimuli, hence their shorter latencies and firing periods (Grill-Spector et al., 2006). Therefore, imagined adaptors enhance the N170/VPP amplitude (compared to perceived adaptors) because they activate neural populations in the visual cortex via top-down processes, thus leaving more bottom-up neural populations ready to fire at full capacity by the time the test stimulus is shown.

Mental imagery and perception also differ in their neural dynamics. For example, Dijkstra et al. (2018) explored the temporal dynamics of category representations during these two processes using magnetoencephalography and multivariate decoding. They found that the visual representations contained in mental imagery become decodable much later after stimulus onset than in perception. In addition, mental imagery exhibited wide temporal generalisation and low temporal specificity compared to perception, which is in line with the view that mental imagery activates visual representations through different top-down connections (Dijkstra et al., 2017; Mechelli et al., 2004). In addition, Canales-Johnson et al. (2021) explored the neural dynamics of mental imagery measuring EEG phase synchronisation. They found that mental imagery of faces entails short-range frontal synchronisation in the theta frequency band and long-range phase synchronisation in the gamma frequency band between frontoparietal and occipitoparietal electrode pairs, in this order. They interpreted these two phase-synchronisation periods as signatures of top-down mnemonic reactivation and endogenous visual binding of facial features, respectively.

How do visual hypnotic hallucinations differ from regular mental images in their top-down processes and temporal neural dynamics? Here we used an adaptation task based on the paradigm developed by Ganis & Schendan (2008) in order to explore how hypnotic hallucinations engage with top-down processes. We measured the N170/VPP complex to face images that were preceded by a hallucinated, imagined, or perceived face.

Subsequently, we used multivariate decoding to explore the neural dynamics of hypnotic hallucination. After testing 130 participants for level of hypnotisability and ability to hallucinate, we selected a group of 16 hypnotic virtuosos to take part in our study, half of whom were able to create vivid visual hallucinations. To our knowledge, our study has recruited the highest number of hypnotic virtuosos to date. By studying hallucinators and non-hallucinators, our findings provide a stringent picture of the neural processing of hypnotic hallucinations. Data and materials are publicly available on the Open Science Framework https://osf.io/p3htg/.

## 2 Methods

### 2.1 Participants

A hundred and thirty university students (all between 18 and 35 years old) attended one of five hypnotisability assessment sessions. We employed the Harvard Group Scale of Hypnotic Susceptibility, Form A (HGSHS:A) to assess their level of hypnotisability. Twenty-seven participants scored 10 or higher out of 12 points (i.e. very highly hypnotisable or hypnotic virtuosos), and were invited to participate in the study. Sixteen hypnotic virtuosos took part in the experiment. Two participants were later excluded from the analysis due to EEG artifacts (see analysis section). The remaining 14 participants (7 female; all right-handed) had a mean age of 22.71 [*SD*_age_ = 3.73]. They were divided into two groups based on whether they passed the hallucinatory task in the HGSHS:A (item 9: “fly hallucination”, in which they are suggested to hallucinate a fly flying around), leaving 7 participants in the hallucinators group (*M*_age_ = 22.14 [3.93]; *M*_HGSHS:A_ = 10.3 [1.38]; 4 female) and 7 participants in the non-hallucinators group (*M*_age_ = 23.29 [3.73]; *M*_HGSHS:A_ = 9.14 [0.378]; 3 female). Finally, we randomly invited 12 moderate hypnotisable participants (4 ≤ HGSHS:A ≤ 8) of whom 10 participated, thus conforming a control group (*M*_age_ = 23 [4.16]; *M*_HGSHS:A_ = 6.1 [1.79]; 6 female). All participants (n = 24; *M*_age_ = 22.83 [3.83]) were right-handed with normal or corrected-to-normal vision, no history of psychiatric or neurological disorders, and no current use of any psychoactive drugs. Participants did not differ in age, gender, or education between groups. All provided informed consent in accordance with the Declaration of Helsinki. This study was approved by the ethics committee of Universidad Diego Portales Faculty of Psychology (Chile).

### 2.2 Stimuli

Stimuli were 70 greyscale face images of highly recognisable celebrities in Chile, 70 greyscale images of highly recognisable objects (selected from the Bank of Standardized Stimuli published by Brodeur et al., 2010), one greyscale image of an oval, and 140 names (the celebrities’ and objects’ names in Helvetica font). All stimuli were equated in luminance and contrast using Adobe® Photoshop® and were presented on a black background. Face images were 7.32 × 8.87 cm in size, front view, with hair removed, and presented inside an oval that subtended ~4.19 × 5.08° of visual angle. Stimuli were presented on a 17-inch LCD monitor using Python and PyGame software. Participants sat 100 cm from the computer monitor and placed their right hand near the spacebar of a keyboard. See Supplementary Material 1 for a description of the stimulus validation procedure.

### 2.3 Procedure

#### 2.3.1 Stimuli study session

The task was a modified version of the adaptation paradigm developed by Ganis & Schendan (2008). Prior to the experimental task, while the EEG cap was being applied, participants had to study each stimulus and its corresponding name. Each image was presented for 300 ms, preceded by its name. Participants were told that later on they would be asked to visualise them from memory. They were also encouraged to study each image for as long as necessary before moving on to the next image. Each stimulus was presented and studied 12 times. Participants had a 10-minute break before continuing to the next condition.

#### 2.3.2 Visual imagery practice

Next, participants had to practise visualising each studied image three times. Each image’s name was presented followed by a grey oval positioned at the middle of the screen. Participants were instructed to visualise each image inside the grey oval and to press a key once their mental image was clear. 200 ms after the keypress, the actual image was presented for 300 ms to help them adjust their mental image. Participants were given a maximum of 10 s to press the key. If no response was received, the task moved on to the next trial. This session had 280 trials, with 2 trials per stimulus – identical stimuli were not presented contiguously.

#### 2.3.3 Hypnotic hallucinatory condition

Only hypnotic virtuosos underwent this condition. Participants underwent a standard (scripted) hypnotic induction procedure performed by a clinical psychologist and cognitive neuroscientist trained in clinical hypnosis (R.C.L.). This procedure involved classic techniques of hypnotic induction and deepening such as suggestions of muscular relaxation, eye fixation, heaviness of the eyelids, arm levitation, etc. After participants exhibited physical signs of hypnotic state such as dropping of the lower jaw, slow breathing rate, facial relaxation, etc., they were administered hypnotic suggestions of visual hallucination (a.k.a. “deceptive suggestions”, see Koivisto et al., 2013) in a direct yet flexible manner, following a “if x then y” structure. Participants were suggested that whenever they saw an image name (celebrities’ or objects’ names) on the screen, they would see the corresponding image inside the grey oval to be presented next. The suggestions also indicated that the resulting visual experience would be vivid, automatic, and felt like it was real. Participants were asked whether they understood the instructions. All participants replied that they did. Next, participants were given hypnotic suggestions of amnesia - they would not remember having heard any of these suggestions even though they would still experience the visual hallucinations as described. Finally, participants had to count from one to ten at their own pace and open their eyes as soon as they wanted to after finishing counting. These hypnotic suggestions of hallucination can be described also as ‘posthypnotic suggestions’ since their effects were indicated to take place after participants had opened their eyes.

Participants were asked to sit comfortably, face the computer monitor, and pay attention to the centre of the screen. Trials began with the name of a celebrity or object presented on the screen for 300 ms. 200 ms later, a grey oval was presented. Participants were instructed to press a key as soon as they saw (i.e. hallucinated) the corresponding (adaptor) image. 200 ms after the keypress an image appeared for 300 ms (test stimulus), which was congruent with the face or object name presented before (Figure 1a).

**Figure 1.**
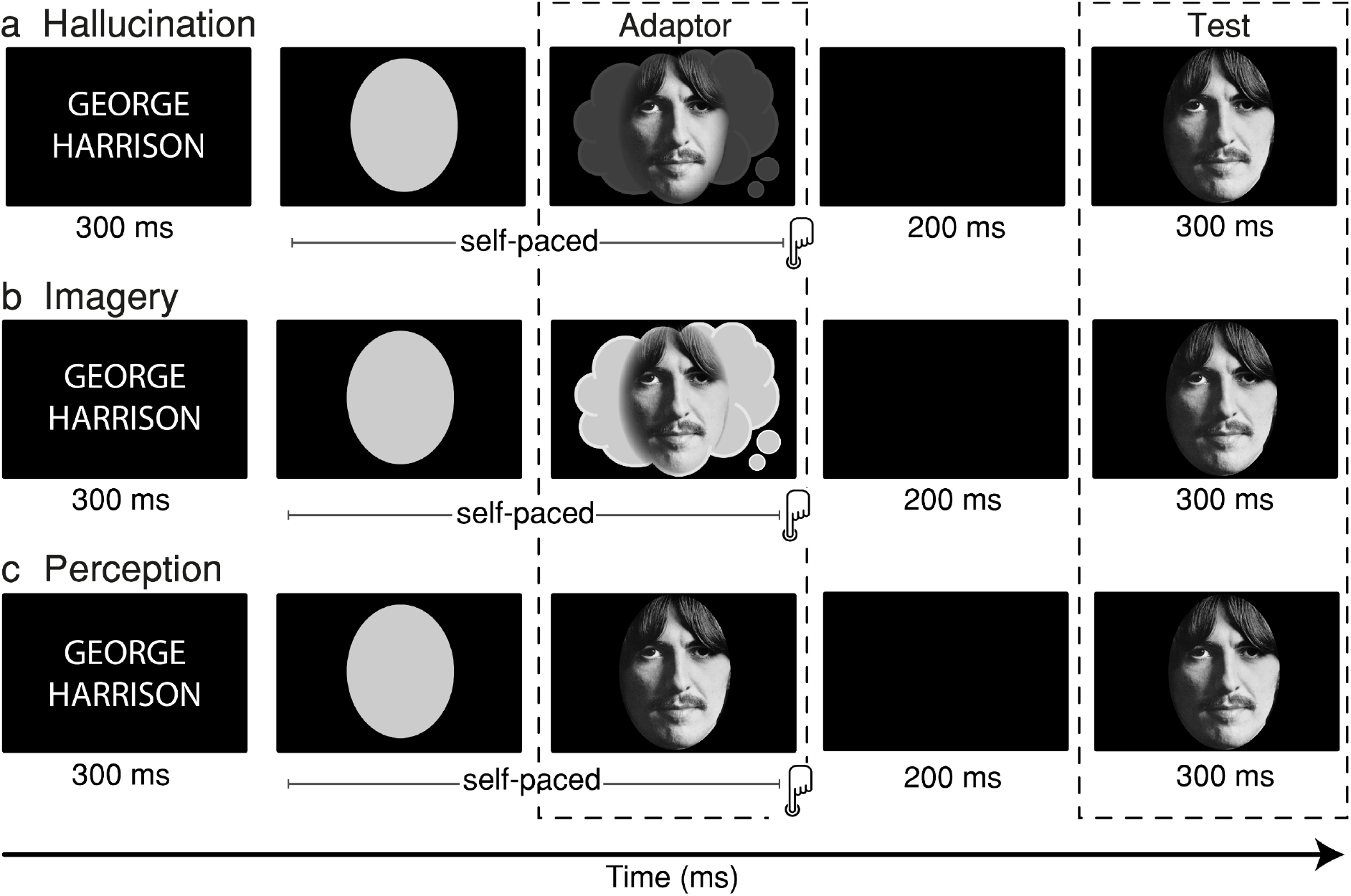
Schematics of a trial. (a) Hallucinatory condition. Before the experiment, participants were instructed under hypnosis (posthypnotic suggestion) that whenever they saw a name on the screen, they would automatically see its corresponding image inside the oval that appeared right after, and that this image would be experienced vividly and as it was real (adaptor stimulus). During the task, they were instructed to indicate with a keypress as soon as they saw an image (self-paced). After the keypress, 200 ms passed before the screen showed the real image (test stimulus) for 300 ms. (b) Imagery condition. Participants were instructed to visualise inside the grey oval the image that corresponded to the name shown at the beginning of the trial and to press the response key as soon as they had a clear mental image. The rest of the trial was as in the hallucinatory condition. (c) Perceptual condition. Participants were instructed to press a key as soon as they saw an image inside the grey oval. After the keypress, 200 ms passed before the screen showed the test stimulus.

There were 280 trials in total, of which half contained a face image as test stimulus whereas the other half contained an object image. Each stimulus was tested twice. Similarly, half of the trials required participants to hallucinate a face and the other half required them to hallucinate an object as adaptor stimulus. The task involved four 70-trial blocks: (1) hallucinated face followed by test face; (2) hallucinated object followed by test face; (3) hallucinated face followed by test object; and (4) hallucinated object followed by test object. Trials were randomised. At the end of this condition, participants were asked to rate the vividness of their (hallucinated) visual experience on a 1-10 Likert scale: *“From 1 to 10, how vivid does the image feel? For example, 1 means it feels like you are imagining it and 10 means it feels like it is really there on the screen”*.

Finally, participants were given suggestions to cancel the hallucinatory effects, and any other hypnosis-related effects (e.g. Koivisto et al., 2013; McGeown et al., 2012; Terhune et al., 2010). Participants had a 10-minute break before continuing to the next condition.

#### 2.3.4 Visual imagery condition

All participants underwent this condition. As shown in Figure 1b, this condition had a very similar structure to the hypnotic hallucinatory condition, with only two differences: firstly, this condition did not involve hypnotic induction or suggestions and, secondly, participants were instructed to visualise each image inside the grey oval (adaptor stimulus) and to press a key once their mental image was clear. 200 ms after the keypress, the actual image was presented for 300 ms (test stimulus). Before continuing to the next condition, participants took another 10-minute break.

#### 2.3.5 Perceptual condition

All participants underwent this condition, which did not involve hypnotic induction or suggestions. Each trial began with the name of a celebrity or object that was shown on the screen for 300 ms (Figure 1c). 200 ms later, a grey oval was presented. Participants were instructed to press a key as soon as they saw a face or object inside the grey oval (adaptor stimulus). 200 ms after the keypress, an image appeared for 300 ms (test stimulus), which was congruent with the face or object name shown before.

### 2.4 EEG recording and pre-processing

EEG data were recorded and digitised using a GES300 Electrical Geodesic amplifier at a sampling rate of 500 Hz and 129-channel saline-based HydroCel sensor nets. Physical filters were set at .01-100 Hz during recording acquisition, and a 50-Hz notch filter was applied offline to remove the DC component. Scalp electrodes were referenced to Cz and impedance values were kept below 50 kΩ. Post-acquisition, the continuous EEG data were resampled to 256 Hz, filtered for frequencies between .3 and 40 Hz, and finally epoched from 200 ms before to 600 ms after participants’ keypress (adaptor stimulus) or test stimulus presentation, depending on the ERP analysis (see below). An independent-component analysis (ICA) was performed on the epoched EEG signal. Components attributed to eye blinks, ocular movements, heartbeat, and channel noise were removed. Trials with voltage fluctuations exceeding ±150 μV were excluded from further analysis. There was no significant difference in the number of rejected trials between conditions (rejected trials < 2% per condition, *p* > .3). Each participant yielded a minimum of 92% of artifact-free trials. Finally, the EEG signal was re-referenced to the average across all electrodes.

### 2.5 Event-related potential analysis

The EEGLAB Matlab toolbox was employed for data pre-processing and pruning (Delorme & Makeig, 2004). Waveforms were averaged for all electrodes and by eye inspection on canonical sites, we determined the following regions of interest (ROI) for each event-related potential (ERP) component of interest: left N170 (65, 69, and 70), right N170 (83, 89, and 90), and VPP (6, 7, and 106). Each triple of electrodes was averaged. Next, mean amplitudes were computed within the 170 - 200 ms time window; this window was determined by eye inspection on the grand average plots restrained to canonical time windows for the N170/VPP complex (Eimer, 2011; Eimer & Holmes, 2007), and did not involve exploratory statistical testing in order to avoid increasing familywise error (Luck, 2014; Luck & Gaspelin, 2017). Two participants (one hallucinator and one non-hallucinator) were excluded from further analysis due to voltage fluctuations exceeding ±200 μV on above 15% of trials.

### 2.6 Source reconstruction analysis

ERP cortical sources were reconstructed using Brainstorm (Tadel et al., 2011; version released in December 2020). To estimate the cortical source of an ERP waveform, we need to model the electromagnetic properties of the head and the sensor array (forward model) and, subsequently, use this model to produce a constrained model of the brain sources that produced the EEG signal of interest (inverse model). The forward model was calculated using the Open MEEG Boundary Element Method (Gramfort et al., 2010) on the cortical surface of a template MNI brain (ICBM152) with 1 mm resolution. The inverse model was constrained using weighted minimum-norm estimation (wMNE; Baillet et al., 2001) to reconstruct source activations in picoampere-meters. wMNE searches for a distribution of sources with the minimum current that can account for the EEG data of interest. Grand-averaged activation values were corrected by subtracting the mean of the baseline period (−200 to 0 ms before stimulus onset) and smoothed with a 5-mm kernel.

### 2.7 Statistical analysis

Reaction times (RTs) and ERP mean values were submitted to repeated-measures Analysis of Variance (ANOVA) models. For the experimental group (7 hallucinators and 7 non-hallucinators, separately), the data were submitted to a 3 (adaptor condition: hallucination, imagery, perception) × 3 (ROI: left, right, central) repeated-measures ANOVA. For the control group (all 24 participants, including moderate hypnotisable and hypnotic virtuosos), the data were submitted to a 2 (adaptor condition: imagery, perception) × 3 (ROI: left, right, central) repeated measures ANOVA. Greenhouse-Geisser adjusted degrees of freedom were reported when Mauchly’s test indicated a violation of the sphericity assumption. All p-values were Bonferroni-corrected for multiple comparisons. For Bayes factor analysis, we defined the null hypothesis as no difference between conditions by using a standard Cauchy prior distribution centred on zero of .707. Frequentist analyses were performed using Matlab (2020a; MathWorks, Inc.) whereas Bayesian analyses were performed using JASP (JASP Team, 2020). The results were later on confirmed using R (R Core Team, 2020).

### 2.8 Multivariate decoding analysis

In order to complement ERP analysis, a multivariate decoding analysis on the raw EEG data was applied using the ADAM toolbox (Fahrenfort et al., 2018). Multivariate decoding analysis is more sensitive than univariate techniques such as ERP analysis as it can detect neural processing patterns that may be too subtle or complex to affect averaged ERP waveforms (Fahrenfort et al., 2018; Grootswagers et al., 2016; Hebart & Baker, 2018). As shown by Xue & Hall (2015), it is essential to keep a balanced number of trials between conditions when performing a multivariate decoding analysis since design imbalances may have unintended effects on the linear discriminant analysis (LDA; the classification algorithm used here) and area under the curve accuracy metric (AUC; the accuracy performance metric used here). To keep a balanced number of trials across conditions, we randomly selected and discarded trials when necessary (“undersampling”; see Fahrenfort et al., 2018). We quantified classifiers’ accuracy performance by measuring the AUC of the receiver operating characteristic (ROC), a measure derived from signal detection theory (Bradley, 1997; Wickens, 2001) that is insensitive to classifier bias. AUC corresponds to the total area covered when plotting the cumulative true positive rates against the cumulative false positive rates for a given classification task (Wickens, 2001). Thus, finding above-chance performance indicates that there was information contained in the neural data that the classifier decoded based on the stimulus features of interest.

We used the whole epoched EEG data time-locked to test stimulus presentation. We classified these epochs according to the nature of their adaptor stimulus (i.e. hallucinated, imagined, or perceived). Next, a backward decoding algorithm using adaptor category as class was applied. We used separate data sets for training and testing; the classifier was trained on the total number of trials across six participants and tested on the remaining participant’s data (leave-one-out cross-validation procedure). This procedure was performed seven times so each participant’s data were left out once for testing. We followed this procedure to maximise the number of trials given the small samples per group. Each iteration provided an AUC score; they were subsequently averaged to obtain a single AUC score per trial time point using t-tests against 50% chance accuracy. These t-tests were double-sided and corrected for multiple comparisons using cluster-based 1000-iteration permutation tests (Maris & Oostenveld, 2007) with a standard cut-off p-value of .05. Finally, as explained below, we also tested whether a paired comparison (“hallucination vs imagery”) was decodable in right-hemisphere electrodes only. For this analysis, electrodes 7, 12, 13, 18, 19, 20, 21, 22, 23, 24, 25, 26, 27, 28, 29, 30,31, 32, 33. 34, 35, 36, 37, 38, 39, 40, 41, 42, 43, 44, 45, 46, 47, 48, 49, 50, 51, 52, 53, 54, 56, 57, 58, 59, 60, 61, 63, 64, 65, 66, 67, 68, 69, 70, 71, 73, 74, 127, and 128 were classified as left hemisphere, and electrodes 1, 2, 3, 4, 5, 8, 9, 10, 14, 76, 77, 78, 79, 80, 82, 83, 84, 85, 86, 87, 88, 89, 90, 91, 92, 93, 94, 95, 96, 97, 98, 99, 100, 101, 102, 103, 104, 105, 106, 107, 108, 109, 110, 111, 112, 113, 114, 115, 116, 117, 118, 119, 120, 121, 122, 123, 124, 125, and 126 as right hemisphere.

## 3 Results

### 3.1 Behavioural results

#### 3.1.1 Reaction times

We explored differences in RTs to adaptor face images. As revealed by a paired-sample t-test, RTs in the perceptual condition (*M*=787 [SD = 491]) were significantly shorter than in the imagery condition (*M*=3390 [1721]) for all 24 participants (*t*(23) = 7.32, *p* < .001, *d* = 1.49), indicating that the visual imagery task took longer than the perceptual task (Figure 2a).

**Figure 2.**
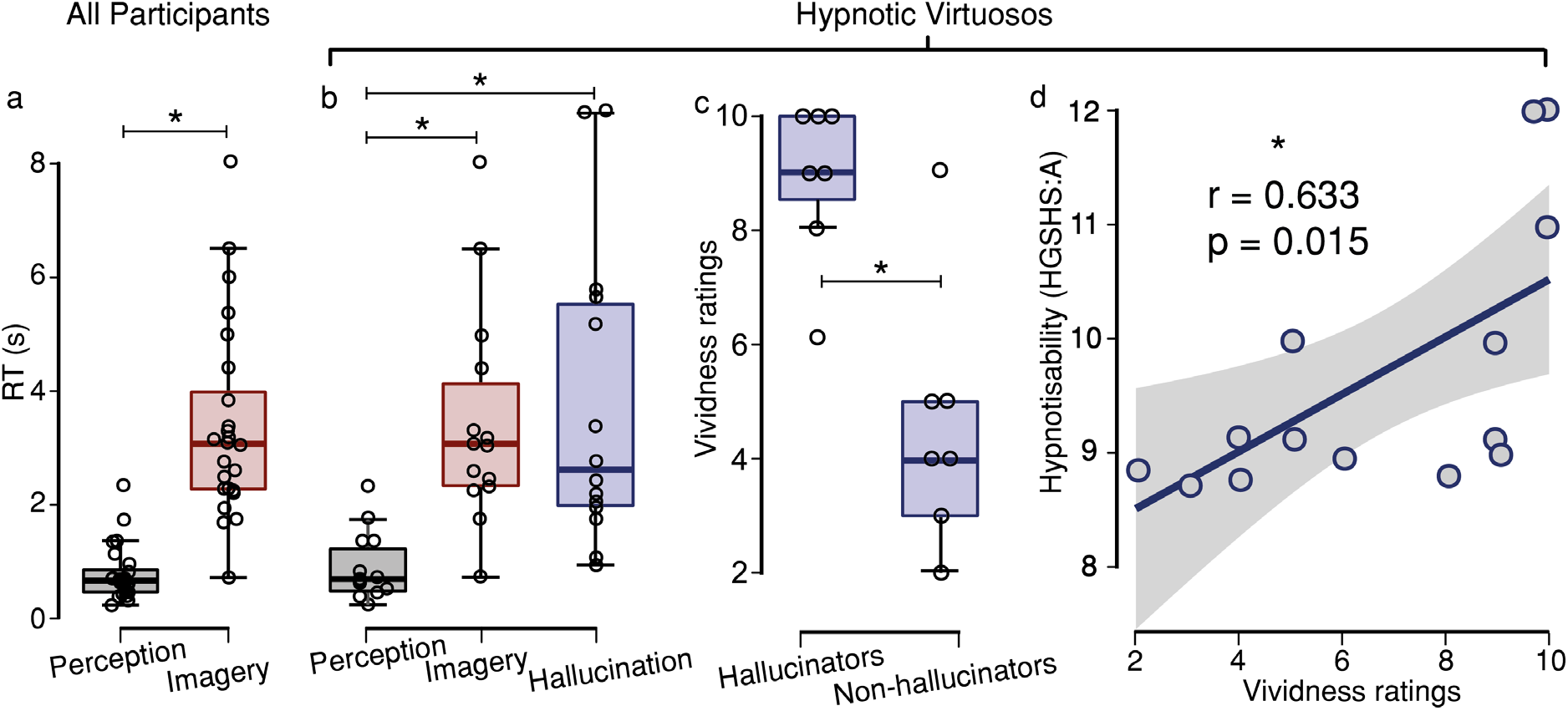
Behavioural results. (a) All participants (n = 24) showed significantly shorter RTs to adaptor faces in the perceptual condition than in the imagery condition. (b) Hypnotic virtuosos (n = 14) showed significantly shorter RTs in the perceptual condition than in the imagery and hallucinatory conditions. RTs during imagery and hallucinatory conditions did not differ significantly. (c) Vividness ratings given by hypnotic virtuosos (n = 14) during the hallucinatory condition. At the end of this condition, participants were asked to rate how vivid the images they visually experienced appeared to them. They had to give a rating in a 1-10 scale. Vividness ratings significantly differed between hallucinators and non-hallucinators. (d) Simple linear regression between hypnotisability (HGSHS:A scores) and vividness ratings. Both variables showed significant positive correlation. Asterisks denote significant effects (*p* < .05).

For the 14 hypnotic virtuosos, we found a main effect of condition on RTs, as revealed by a mixed ANOVA (*F*_(1.49, 17.86)_ = 8.79, *p* = .004, ηp^2^ = .423), which included-hallucination (*M*=3789 [2676.87]), imagery (*M*=3476.21 [1940.19]), and perception (*M*=881.86 [601.6]) as conditions, and group (hallucinators and non-hallucinators) as between-subject factor. Bonferroni-corrected pairwise comparisons found that RTs during perception were significantly shorter than during imagery (*t* = −5, p < .001, *d* = −1.337) and hallucination (*t* = −4, *p* = .004, *d* = −1.071), whereas no difference was found between imagery and hallucinatory conditions (*t* = −0.346, *p* = 1, *d* = −0.093). There was no effect of group (*F*_(1.49, 17.86)_ = 0.059, *p* = .897, ηp^2^ = .005). Finally, we estimated Bayes factors to assess the null effect between hallucinatory and imagery conditions, which provided moderate support for the null hypothesis (*BF*_01_ = 3.514), (Albert, 2009; Ortega & Navarrete, 2017). These results indicate that both the imagery and hallucinatory tasks required longer time than the perceptual task (Figure 2b).

#### 3.1.2 Vividness ratings and hypnotisability scores in hypnotic virtuosos

Not all hypnotic virtuosos exhibited the ability to hallucinate, as assessed by item 9 in the HGSHS:A (see Methods section). Nevertheless, both groups (hallucinators and non-hallucinators) underwent the hallucinatory condition. To explore their visual hallucinatory experience further, we asked them to rate the vividness of their visual experience (i.e. of the adaptor images in the hallucinatory condition). An independent-sample t-test revealed a significant difference in hallucinatory vividness scores (*t*_(12)_ = 4.26, *p* = .001, d = 2.28), with vividness rated higher by hallucinators (*M*=8.86 [1.46]) than non-hallucinators (*M*=4.57 [2.23]; Figure 2c).

We also tested whether they differed in hypnotisability. An independent-sample t-test did not find differences in hypnotisability between hallucinators and non-hallucinators (*t*_(12)_ = 1.47, *p* = .167,d = 7.86). To test whether the evidence supports the null hypothesis, we estimated Bayes factors, which provided strong support for the alternative hypothesis (*BF*_01_ = 0.038;*BF*_10_ = 26.5). These results justified the distinction between hallucinators and non-hallucinators used in the following analyses.

We further tested, in an exploratory manner, the statistical relationship between vividness ratings provided in the hallucinatory condition and hypnotisability scores obtained by the HGSHS:A (Figure 2d). A simple linear regression model showed a positive linear relationship between these variables (*β* = 0.251, *R*^2^ = 0.4, *F*_(1,12)_ = 8.01, *p* = .015), i.e. as hypnotisability increased, so did hypnotic hallucinations’ vividness.

### 3.2 ERP results

#### 3.2.1 All participants: imagery and perceptual conditions

We measured the ERP waveforms elicited by adaptor (first image, either imagined or perceived) and test stimuli (second image, always perceived) in the imagery and perceptual conditions. All participants were included in these analyses, which had two objectives: Firstly, to confirm that the N170/VPP was more sensitive to adaptor faces than adaptor objects, as a sanity check for our method. Secondly, that the imagery and perceptual conditions replicated the adaptation effects on test face stimuli reported by Ganis & Schendan (2008), i.e. that N170 in response to a test face adapted by an imagined face shows significantly greater amplitude than in response to a test face adapted by a perceived face.

##### 3.2.1.1 ERP component evoked by adaptor stimuli

N170/VPP voltage values were significantly more negative to adaptor faces than adaptor objects (*F*_(1, 23)_ = 9.104, *p* = .006, ηp^2^ = .284), thus indicating that our face processing paradigm works. See Supplementary Material 2 for a full description of the N170/VPP evoked by the adaptor stimuli only, i.e. time-locked to the keypress, not to test stimuli.

##### 3.2.1.2 ERP component evoked by test stimuli

###### P1 component

A 2 (adaptor category: face, object) × 2 (condition: imagery, perception) repeated-measures ANOVA was calculated on P1 voltage values evoked by face test stimuli. No effect or interaction reached significance. A Bayes factor analysis provided anecdotal support for the null effect of adaptor category (*BF*_01_ = 2.468) and moderate support for the null effect of condition (*BF*_01_ = 4.399). These results suggest that there were no differences in visual attention between conditions (Hillyard & Anllo-Vento, 1998; Luck et al., 1994).

###### N170/VPP complex

The N170/VPP results replicated the findings reported by Ganis & Schendan (2008), thereby validating the adaptation task. We submitted the data to a 2 (adaptor category: face, object) × 2 (condition: imagery, perception) × 3 (ROI: left N170, right N170, VPP) repeated-measures ANOVA. We found a main effect of condition (*F*_(1, 23)_ = 7.306, *p* = .013, ηp^2^ = .241), a main effect of ROI (*F*_(1.307, 30.057)_ = 27.281, *p* < .001, ηp^2^ = .543), and a significant interaction between condition and ROI (*F*_(1.385,31.855)_ = 34.058, *p* < .001, ηp^2^ = .597). Bonferroni-corrected pairwise comparisons revealed that imagery had more negative voltage values than perception at left (*t* = 4.593, *p* < .001) and right N170 (*t* = 5.251, *p* < .001) whereas the opposite direction was found at VPP (*t* = −5.736, *p* < .001).

Additionally, the interaction between adaptor stimulus and condition was also significant (*F*_(1, 23)_ = 5.369, *p* = .03, ηp^2^ = .189). Bonferroni-corrected pairwise comparisons per ROI showed that voltage values evoked by test faces that were adapted by imagined faces were significantly more negative than test faces adapted by perceived faces at left (*t* = 3.652, *p* = .028) and right N170 (*t* = 4.668, *p* < .001), and, as expected, the effect was found in the opposite direction at VPP (*t* = −5.934, *p* < .001), (Figure 3) - this replicates the main adaptation effect reported by Ganis & Schendan (2008).

**Figure 3.**
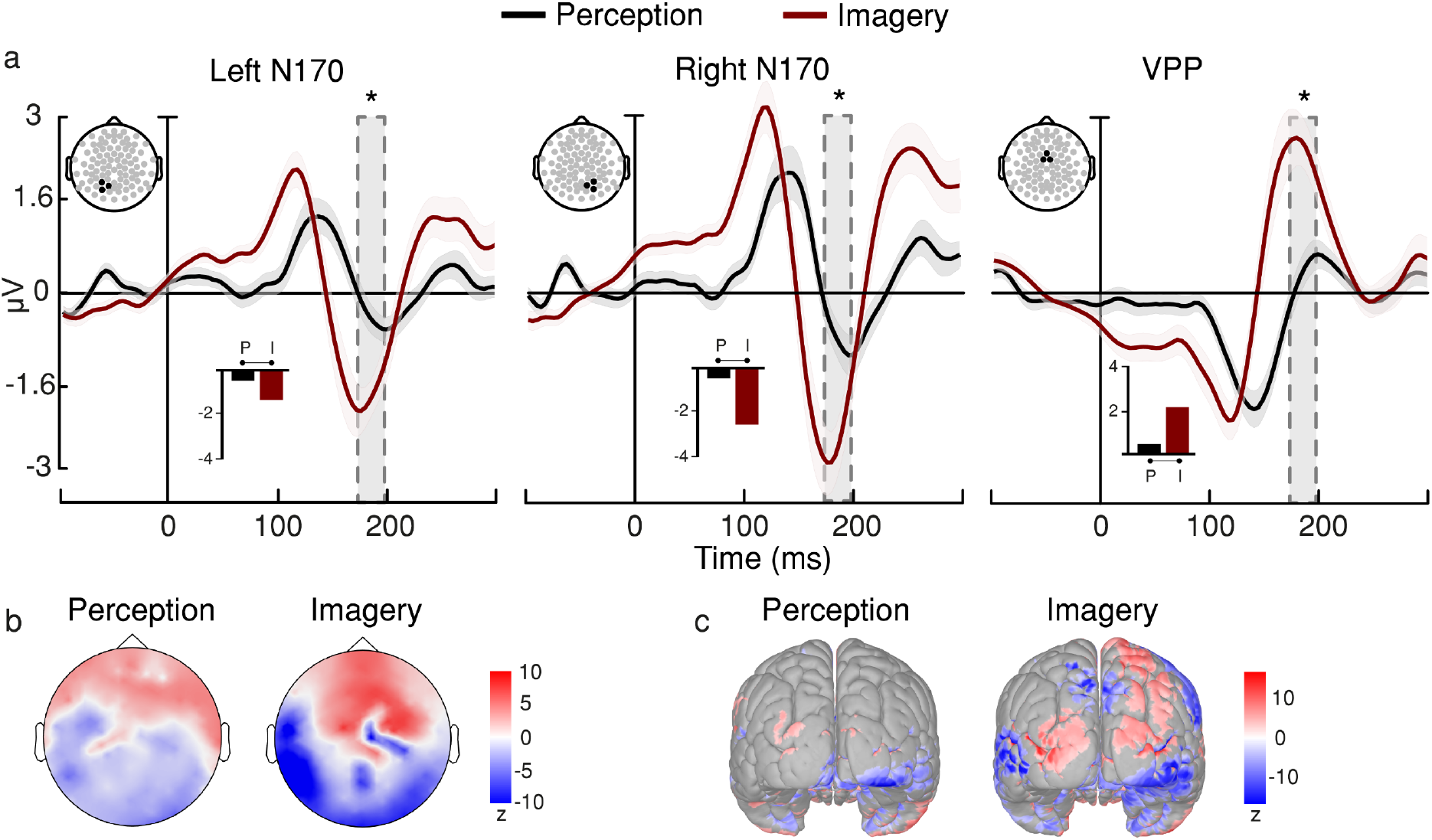
All Participants. N170/VPP evoked by test faces. (a) At left and right N170, a significantly less negative voltage was found in the perceptual condition compared with the imagery condition, indicating a suppression effect of the perceived adaptor faces on test faces. The same effect was found at VPP but in the opposite direction. Asterisks denote significant differences (*p* < .05) between perceptual and imagery conditions. Shaded areas represent standard error of the mean (SEM). Bar plots summarise mean differences. (b) Topographic maps represent voltage distributions in z-scores across the scalp for the time window of interest. (c) Source estimation of N170/VPP at its peak. Variations of current are represented in z-scores.

#### 3.2.2 Hallucinators: hallucinatory, imagery, and perceptual conditions

Does hypnotic hallucination of faces adapt the N170/VPP complex differently than mental imagery? To test our main hypothesis, we submitted the data to a 3 (condition: hallucination, imagery, perception) × 3 (ROI: left N170, right N170, VPP) repeated-measures ANOVA (Figure 4), only including trials with faces as both adaptor and test stimuli (e.g. see Figure 1). We found main effects of condition (*F*_(1.56, 9.358)_ = 10.532, *p* = .006, ηp^2^ = .637) and ROI (*F*_(1.904, 11.425)_ = 34.314, *p* < .001, ηp^2^ = .851). The interaction also reached significance (*F*_(2.299, 13.794)_ = 31.39, *p* < .001,ηp^2^ = .84). Bonferroni-corrected pairwise comparisons revealed that voltage values in the hallucinatory condition (*M*=−2.265 [1.324]) were more negative than in the perceptual condition at left N170 (*M*=−0.132 [0.705]), (*t* = −4.949, *p* < .001). Similarly, voltage values in the imagery condition (*M*=−1.819 [1.039]) were more negative than in the perceptual condition at the same ROI (*t* = −3.915, *p* = .014). However, hallucination and imagery conditions did not differ at left N170 (*t* = −1.034, *p* > .1). Crucially, however, we found that voltage values in the hallucinatory condition (*M*=−4.166 [0.742]) were significantly more negative than in the imagery condition at the right N170 (*M*=−2.623 [1.136]), (*t* = −3.58, *p* = .036), and that the voltage values in the imagery condition were significantly more negative than in the perceptual condition (*M*=−0.415 [1.045]), (*t* = −5.125, *p* < .001). Therefore, we found that hypnotic hallucination differs from mental imagery in how it modulates the N170/VPP complex only in the right hemisphere’s N170 component - this is our main finding. See Supplementary Material 3 for a detailed statistical account.

**Figure 4.**
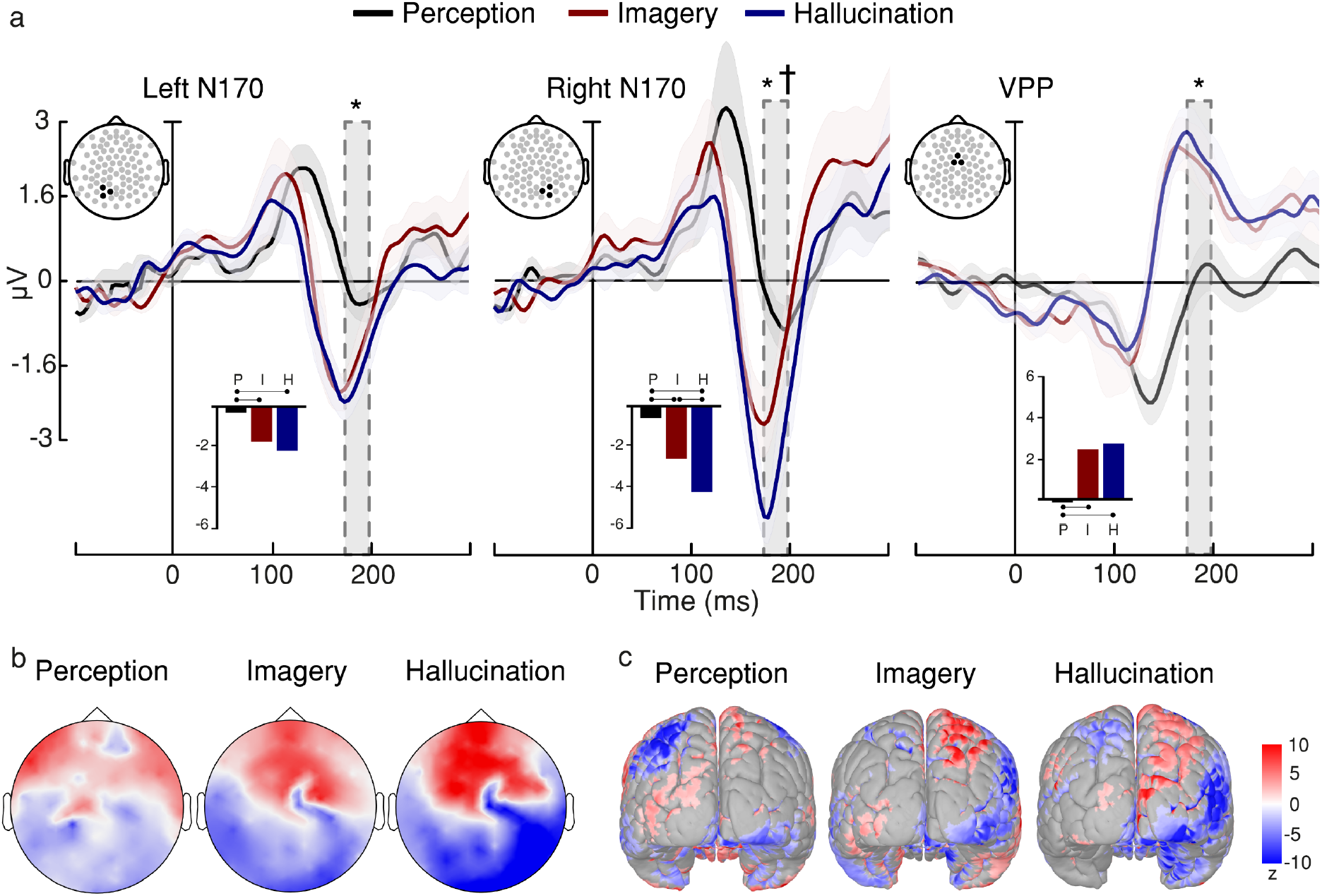
Hallucinators group. N170/VPP evoked by test faces preceded by adaptor faces. (a) At left and right N170, a significantly less negative voltage was found in the perceptual condition compared with the hallucinatory and imagery conditions. The same effect was found at VPP but in the opposite direction. Crucially, we found a lateralised effect at the right N170: significantly more negative voltage values were found in the hallucinatory condition compared with the imagery condition. Shaded areas represent SEM. Bar plots summarise mean differences. Asterisks denote significant differences (*p* < .05) between perceptual and imagery conditions, and between perceptual and hallucinatory conditions. Daggers denote significant differences between hallucinatory and imagery conditions. (b) Topographic maps represent de voltage distributions in z-scores across the scalp for the time window of interest. (c) Source estimation of N170/VPP at its peak. Variations of current are represented in z-scores.

#### 3.2.3 Non-hallucinators: hallucinatory, imagery, and perceptual conditions

Do hypnotic virtuosos who do not hallucinate show this lateralised effect of hallucination on right N170? We entered the data into a 3 (condition: hallucination, imagery, perception) × 3 (ROI: left N170, right N170, VPP) repeated-measures ANOVA (Figure 5), only including trials with faces as both adaptor and test stimuli (e.g. see Figure 1). We found a main effect of condition (*F*_(1.323, 7.941)_ = 5.224, *p* = .045, ηp^2^ = .465) and ROI (*F*_(1.1 79, 7.075)_ — 8.388, *p* = .02,ηp^2^ = .583). The interaction also reached significance (*F*_(2.413, 14.48)_ = 14.532, *p* < .001,ηp^2^ = .708). Bonferroni-corrected pairwise comparisons revealed that voltage values in the hallucinatory condition (*M*=−2.336 [1.322]) were more negative than in the perceptual condition at left N170 (*M*=−0.358 [1.305]), (*t* = −4.171, *p* = .007). Similarly, voltage values in the imagery condition (*M*=−2.277 [1.566]) were more negative than in the perceptual condition at the same ROI (*t* = −4.047, *p* = .001). However, hallucinatory and imagery conditions did not differ significantly at left N170 (*t* = −0.124, *p* > .1). We found equivalent results at right N170: voltage values in the hallucinatory condition (*M*=−2.588 [2.587]) were more negative than in the perceptual condition (*M*=−0.753 [1.852]), (*t* = −3.871, *p* = .017). Similarly, voltage values in the imagery condition (*M*=−2.988 [1.964]) were more negative than in the perceptual condition at right N170 (*t* = −4.714, *p* = .001). Notably, however, we did not find a significant difference between hallucinatory and imagery conditions at right N170 (*t* = 0.843, *p* > .1). Therefore, unlike hallucinators, non-hallucinators did not exhibit such a difference at right N170. Finally, we estimated Bayes factors to assess this null effect between hallucinatory and imagery conditions at right N170, which provided anecdotal evidence in favour of the null hypothesis (*BF*_01_ = 2.274). See Supplementary Material 4 for a detailed statistical account.

**Figure 5.**
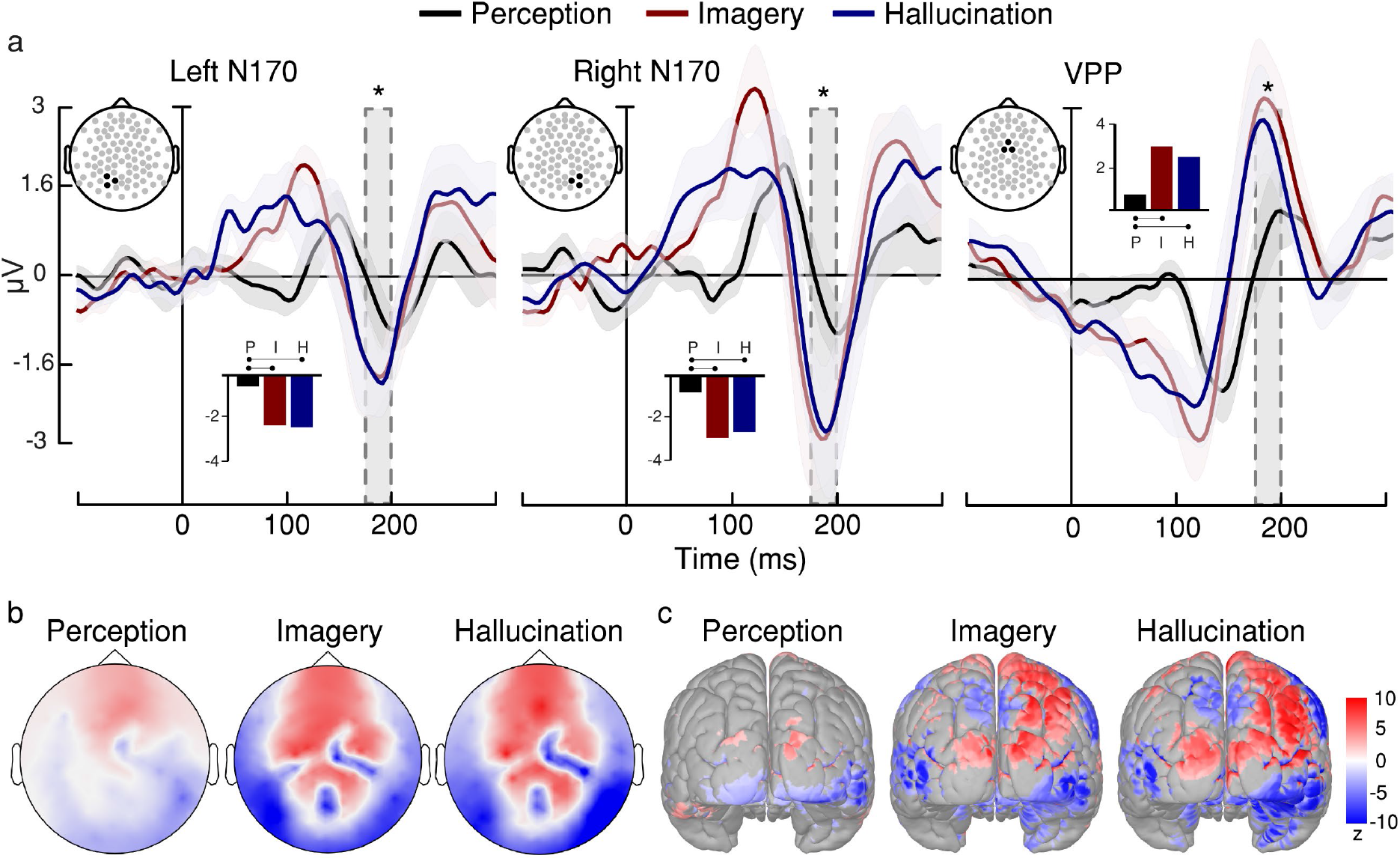
Non-hallucinators group. N170/VPP evoked by test faces preceded by adaptor faces. (a) At left and right N170, a significantly less negative voltage was found in perceptual condition compared with imagery and hallucinatory conditions. The same effect was found at VPP, but in the opposite direction. We did not find a significant difference between hallucinatory and imagery conditions. Asterisks denote significant differences (*p* < .05) between perceptual and imagery conditions, and between perceptual and hallucinatory conditions. Shaded areas represent SEM. Bar plots summarise mean differences. (b) Topographic maps represent de voltage distributions in z-scores across the scalp for the time window of interest. (c) Source estimation of N170/VPP. Variations of current are represented in z-scores.

#### 3.2.4 Group analysis: hallucinatory and imagery conditions

To directly test whether the difference between hypnotic hallucination and mental imagery in right N170 is present in hallucinators but absent in non-hallucinators, and thus ascertain that our findings are due to hypnotic hallucination-related processes, we entered the data into a 2 (condition: hallucination, imagery) × 3 (ROI: left N170, right N170, VPP) × 2 (group: hallucinators, non-hallucinators) mixed ANOVA, only including trials with faces as both adaptor and test stimuli. While we did not find an effect of group (*F*_(1, 12)_ = 0.323, *p* = .58,ηp^2^ = .026), we did find a significant interaction between group and condition (*F*_(1, 12)_ = 4.959, *p* = .046, ηp^2^ = .292). Bonferroni-corrected pairwise comparisons confirmed the existence of a significant difference between hallucinatory and imagery conditions in right N170 for the hallucinators group (*t* = −3.87, *p* = .032), which was not found for the non-hallucinators group (*t* = 1.003, *p* = 1).

Taken together, these results suggest that the ability to hallucinate in response to hypnotic suggestions is associated with a lateralised effect on right N170. In other words, while hallucinators exhibit a modulation difference at right N170 between hallucinatory and imagery conditions, non-hallucinators do not exhibit this modulation.

#### 3.2.5 Relationship between hallucination vividness and N170

To further study the relationship between hypnotic hallucination and N170 adapted evoked response, we performed a multiple regression analysis using left N170 and right N170 voltage values as predictors and vividness ratings as outcome variable. The model predicted the vividness ratings provided in the hallucinatory condition (*R*^2^ = 0.424, *F*_(2,11)_ = 4.05, *p* = .048); right N170 was a reliable predictor (*β* = −0.941, *p* = .016) whereas left N170 was not (*β* = 0.322, *p* = .55), (Figure 6a). These relationships were also described in terms of simple Pearson’s correlations, which showed a significant association between vividness ratings and right N170 voltage values (*r* = −.636, *p* = .015) but not between the former and left N170 voltage values (*r* = .042, *p* = .886), (Figure 6b). These results support our ERP findings in respect to a lateralised effect of hypnotic hallucination.

**Figure 6.**
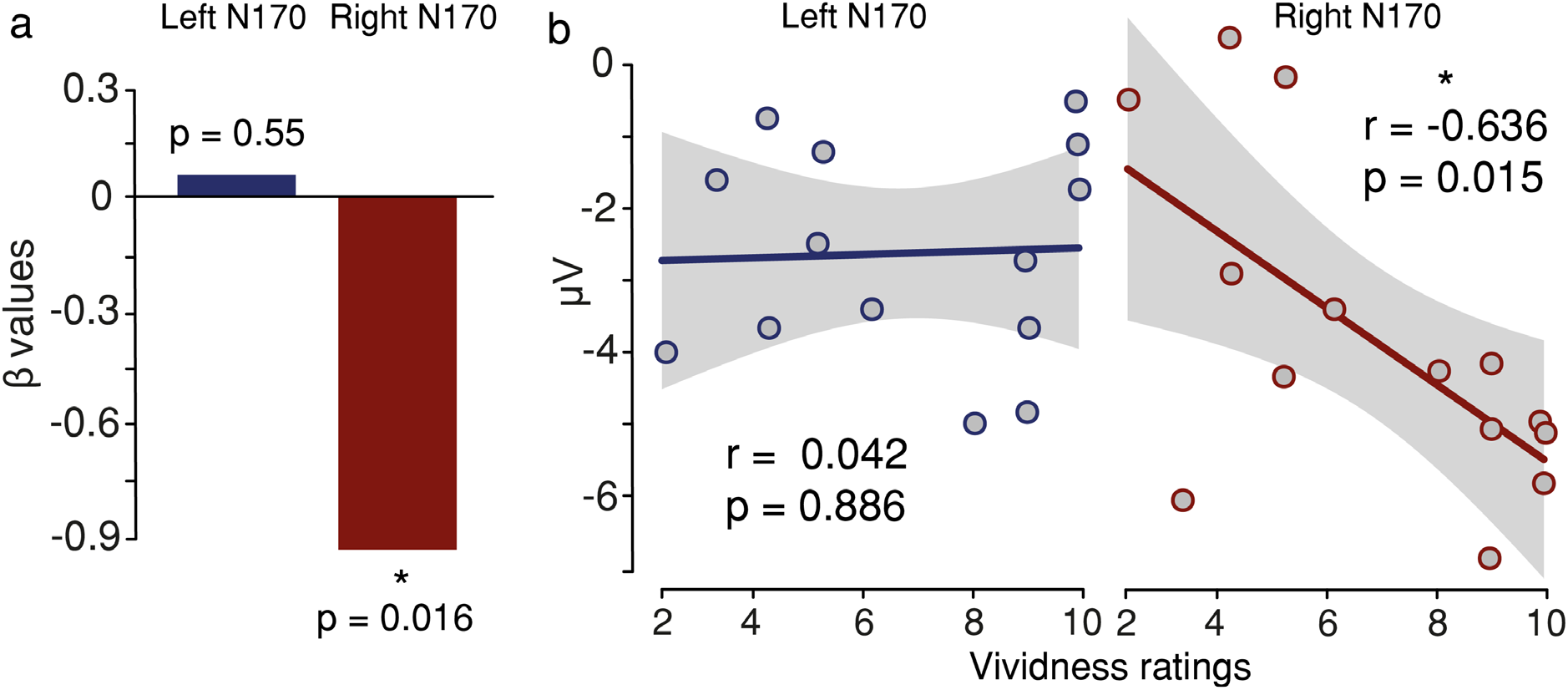
Relationship between vividness ratings and N170 voltage values in hallucinatory condition. (a) Multiple linear regression using left and right N170 voltage values as predictors and vividness ratings as outcome variable. β values for each predictor. Only right N170 was a reliable predictor. (b) Simple regressions. Only vividness ratings and right N170 voltage values showed a significant association - the more vivid the hallucinations, the more negative the voltage values in right N170.

### 3.3 Multivariate decoding results

Given its univariate nature, ERP analysis may not be sensitive to all differences in neural patterns between conditions. To test whether more spatially or temporally extended patterns of neural activity may distinguish these conditions, we performed a multivariate decoding analysis on the raw EEG data. Decoding analyses search for multivariate differences across electrodes without requiring a priori specifications such as electrodes or temporal windows of interest (Fahrenfort et al., 2018). To achieve this, we trained classifiers to: (1) discriminate between hallucinatory, imagery, and perceptual conditions (multiclass decoding) in the adapted EEG signals of interest (i.e. time-locked to test faces preceded by adaptor faces); (2) discriminate between each paired comparison (i.e. paired decoding: ‘hallucination vs imagery’, ‘hallucination vs perception’, and ‘imagery vs perception’); and (3) discriminate ‘hallucination vs imagery’ on left-hemisphere electrodes and right-hemisphere electrodes, separately, to test whether the lateralised effect of hallucination at right-hemisphere N170 reported above replicates using this technique.

#### 3.3.1 Multiclass decoding

Consistent with our ERP findings, multivariate decoding showed that the three conditions were decodable above chance (*p* < .05, cluster corrected) in the hallucinators group, indicating that the information contained in the neural data was processed differently between conditions around the temporal window of the N170/VPP complex (Figure 7a). The temporal generalisation matrix of classification accuracy between training and testing time points shows low temporal generalisation of the decoded patterns, mainly circumscribed to the same time window of the N170/VPP complex (Figure 7b), which may suggest that differences in top-down processing between conditions, as measured by our adaptation paradigm, are temporally restricted. This is in line with our ERP findings and supports our claim that hallucination, imagery, and perception engage with top-down mechanisms differently.

**Figure 7.**
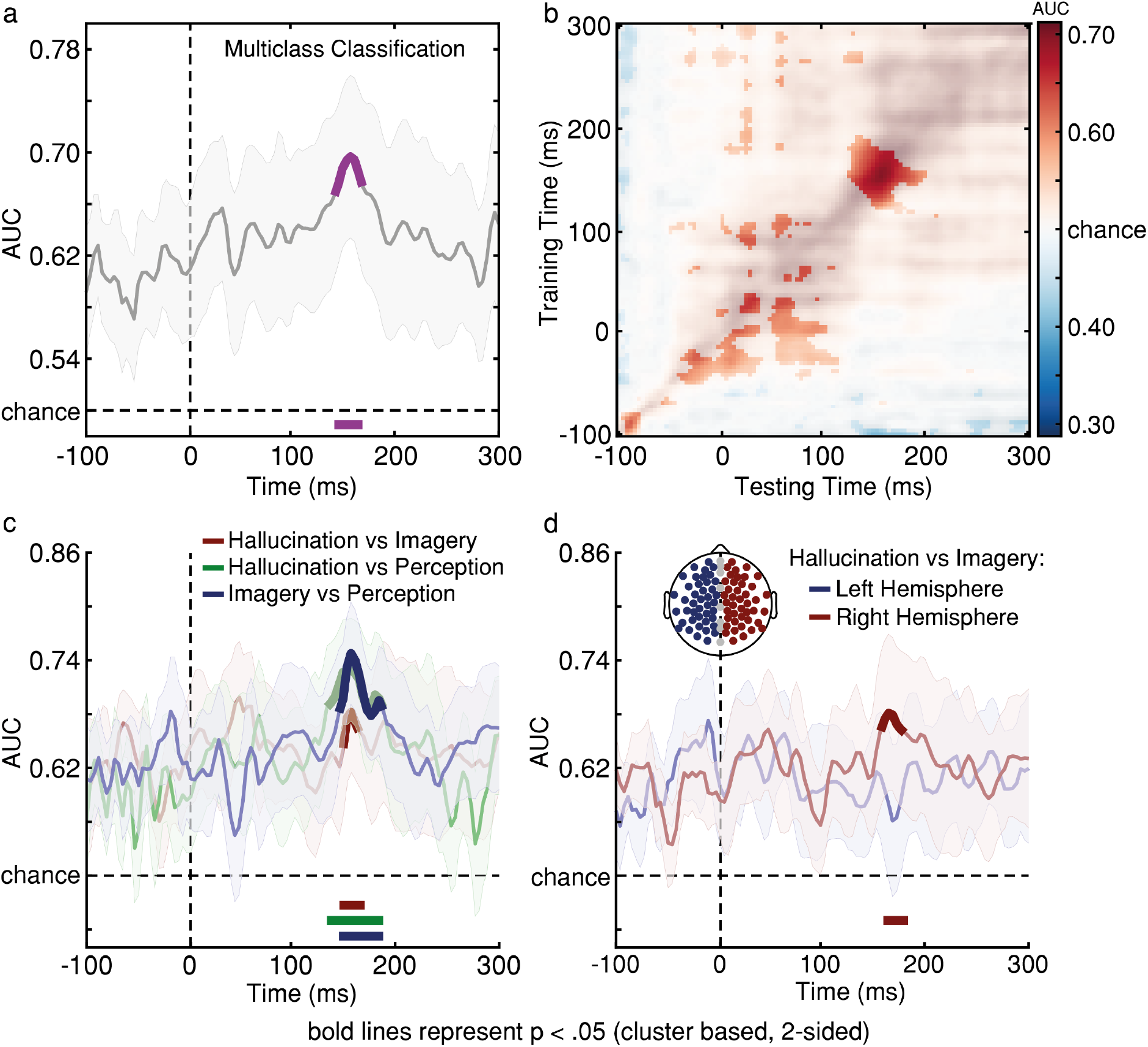
Multivariate decoding analysis in the hallucinators group. (a) Multiclass decoding analysis of hallucination, imagery, and perception conditions. Classifier AUC scores were significantly above chance in the 140-1 75 ms time window. (b) Temporal generalisation matrix of multiclass classification accuracies. The Y-axis depicts when the classifier was trained and the X-axis depicts when it was tested, relative to the test stimulus onset (time zero). (c) Multivariate decoding of paired comparisons: ‘hallucination vs imagery’, ‘hallucination vs perception’, and ‘imagery vs perception’. AUC scores were significantly above chance around the same time window of the N170/VPP complex for all paired comparisons. (d) Multivariate decoding of the ‘hallucination vs imagery’ comparison for left-hemisphere and right-hemisphere electrodes, separately. AUC scores were significantly above chance only in the analysis that included right-hemisphere electrodes. Solid bold lines at the bottom of the chart represent significant clusters (*p* < .05, cluster corrected) and shaded contours represent SEM.

#### 3.3.2 Paired decoding

Are there differences in top-down neural processing specifically between hallucination and imagery, as indicated by our ERP results in the hallucinators group? Multivariate decoding analysis showed that each paired comparison was decodable above chance (*p* < .05, cluster corrected) around the same temporal window shown relevant in the multiclass decoding analysis. Critically, the classifier performed above chance decoding for all paired comparisons, including the ‘hallucination vs imagery’ comparison, although with lower AUC scores than the other two (Figure 7c). As reported above, ERP results showed a lateralised effect on N170; more specifically, only the right-hemisphere N170 showed a significant voltage difference between hallucinatory and imagery conditions. To test for this lateralised effect here, we decoded the comparison ‘hallucination vs imagery’ on left-hemisphere and right-hemisphere electrodes separately. Crucially, we found above-chance decoding performance only when the classifier was trained and tested on right-hemisphere electrodes (Figure 7d), thus supporting a lateralised effect of hallucination over imagery.

In the non-hallucinators group, on the contrary, multivariate decoding analysis could not decode the paired comparison ‘hallucination vs imagery’ with above-chance performance (Figure 8a), regardless of the hemisphere (Figure 8b).

**Figure 8.**
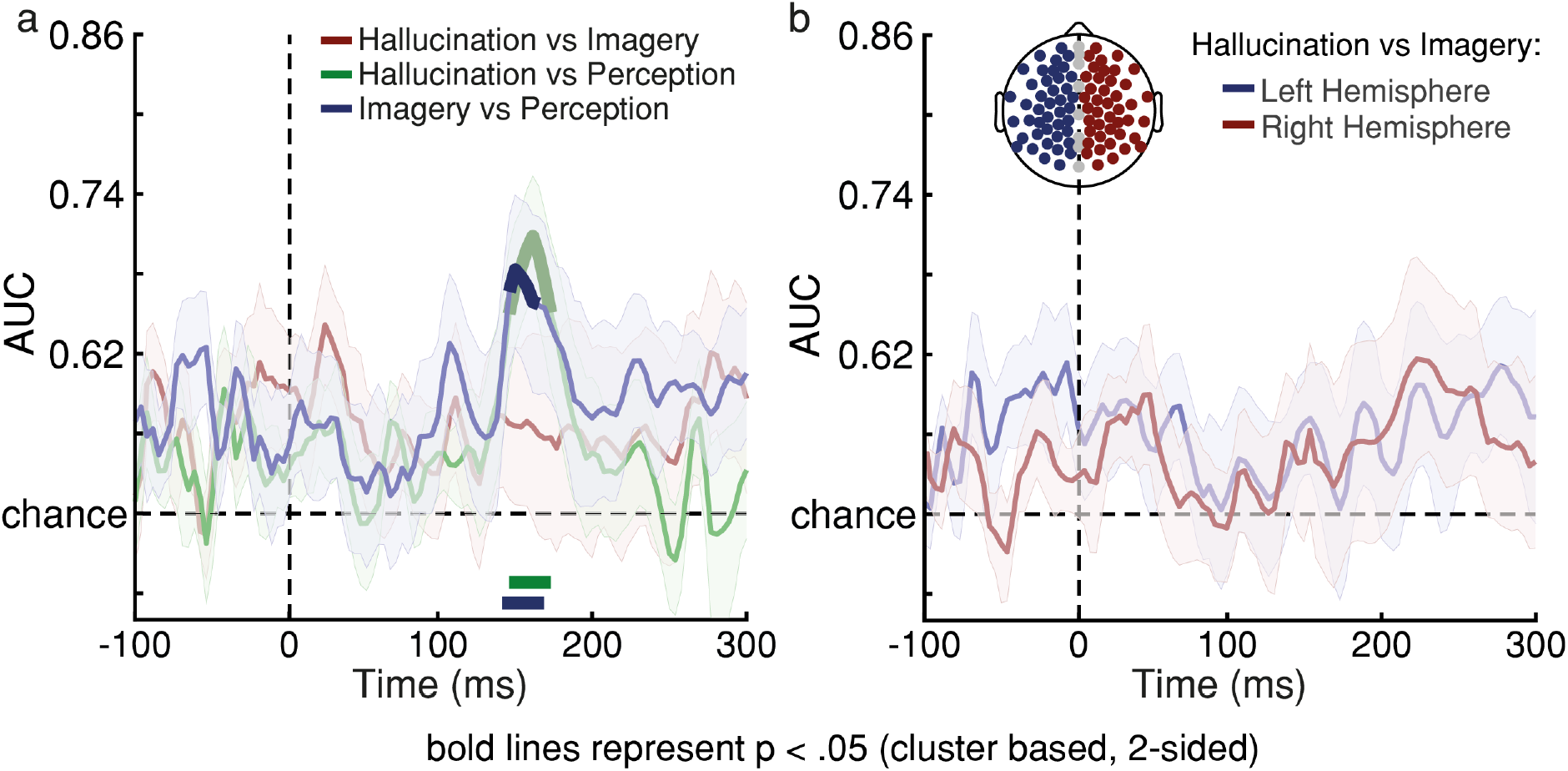
Multivariate decoding analysis in the non-hallucinators group. (a) Multivariate decoding of paired comparisons: ‘hallucination vs imagery’, ‘hallucination vs perception’, and ‘imagery vs perception’. AUC scores were significantly above chance around the same time window of the N170/VPP complex for ‘hallucination vs perception’ and ‘imagery vs perception’ comparisons. (b) Multivariate decoding of the ‘hallucination vs imagery’ comparison for left-hemisphere and right-hemisphere electrodes, separately. The comparison could not be decoded above chance in either group of electrodes. Solid bold lines at the bottom of the chart represent significant clusters (*p* < .05, cluster corrected) and shaded contours represent SEM.

Altogether, ERP results and multivariate decoding results suggest that the neural signature that distinguishes between hypnotic hallucination and mental imagery, at least in relation to top-down modulation over the visual cortex, lies in the right hemisphere.

## 4 Discussion

Does hypnotic hallucination differ from mental imagery in how it endogenously reactivates neural representations of faces? Fourteen hypnotic virtuosos (along with 10 moderately hypnotisable control participants) took part in our study. Based on the hypnotic virtuosos’ performance in a hypnotic hallucinatory task, we categorised them as either hallucinators or non-hallucinators. To date, our study has the largest sample of hypnotic virtuosos among hypnotic hallucination studies (e.g. Kallio & Koivisto, 2013, 2016; Koivisto et al., 2013; Kosslyn et al., 2000); furthermore, by assessing hypnotic virtuosos’ ability to hallucinate, we are able to draw conclusions that are specific to the nature of hypnotic hallucination regardless of hypnotisability differences. We used an adaptation paradigm developed by Ganis & Schendan (2008) to assess top-down reactivation of neural representations during hypnotic hallucination and mental imagery of faces. We measured the adaptation effect of hallucinated, imagined, and perceived faces on the N170/VPP complex and found evidence of increased top-down reactivation in hallucinated faces over imagined faces in the right occipitotemporal cortex. This difference in top-down reactivation increase between hypnotic hallucination and mental imagery at right N170 was only found in the hallucinators group. Since both hallucinators and non-hallucinators were hypnotic virtuosos and did not differ in the hypnotic induction procedure, this effect cannot be attributed to differences in those factors and therefore it may be specific to the neural processing that characterises hypnotic hallucination. Consistent with this, we found a significant correlation between vividness ratings and right N170 voltage values but not between the former and left N170 voltage values. Our multivariate decoding analysis supported these findings: the paired comparison ‘hallucination vs imagery’ was decodable around the same time window of the N170/VPP complex and across right-hemisphere electrodes but not across left-hemisphere electrodes. Altogether, our findings suggest that hypnotic hallucination of faces modulates the visual cortex through lateralised top-down mechanisms in the right hemisphere.

Past studies have speculated about a specialised role of the right hemisphere for hypnosis. For example, Bakan (1969) reported that highly hypnotisable individuals show more reflective eye movements to the left than lowly hypnotisable ones, a finding that was later supported and extended by Gur & Gur (1974). Similarly, Hass & Holden (1987) reported that hypnotised participants exhibited faster visual detection on stimuli shown to the left visual fields than controls (also see Naish, 2010). Several EEG studies found results consistent with this. For example, Edmonston & Moscovitz (1990) found a shift from left-to right-hemisphere activation during hypnosis and Macleod-Morgan & Lack (1982) found the same evoked pattern in highly hypnotisable individuals, especially while engaged in a continuous spatial orientation task. Furthermore, De Pascalis (1993) found greater gamma-band amplitude in the right hemisphere (compared to the left hemisphere, and especially in posterior areas) in highly hypnotisable individuals who were experiencing hypnotically suggested dreams. Studies employing other methods have supported this lateralised effect. For example, using fMRI, Crawford et al. (1983) found an increase in blood flow in the right hemisphere followed by a hypnotic induction procedure, and so did Kosslyn et al. (2000) after hypnotising and suggesting highly hypnotisable individuals to hallucinate colours. Similarly, but measuring electrodermal response, Gruzelier et al. (1984) found that highly hypnotisable individuals that undergo hypnotic induction show a reduction in electrodermal response in their left hand compared to their right hand. Lowly hypnotisable individuals did not show this asymmetry. However, many studies have failed at finding a lateralised effect of hypnosis or hypnotisability (e.g. Graffin et al., 1995; Kihlstrom et al., 2013; Ray, 1997; and also see Jasiukaitis et al., 1996, and Maquet et al., 1999), which has led researchers to gradually abandon the hypothesis of a hemispheric specialisation for hypnosis or hypnotic suggestion (for a critical review, see Kihlstrom, 2013).

Our findings revive the debate about a lateralised neural mechanism for hypnotic suggestion by providing new evidence of a crucial role of the right hemisphere during hypnotic hallucination. Having found a lateralised adaptation effect in hypnotic virtuosos who can hallucinate but not in hypnotic virtuosos who do not suggests that this effect is due to the perceptual changes that the induced hallucinatory experience entails. Our adaptation task tests to what extent an adaptor stimulus relies on top-down processing to activate neural representations of faces. As argued by Ganis & Schendan (2008), perceived adaptors suppress the amplitude of the N170/VPP complex because they affect the neural populations in the visual cortex that support early perceptual processing via bottom-up mechanisms. Since mental imagery activates these early perceptual mechanisms via top-down processes, these neural populations’ firing rate is not suppressed, hence the higher amplitude of the N170/VPP. Following this line of reasoning, hallucinated adaptors must have activated neural representations of faces via top-down mechanisms to a greater extent than mental imagery did. The fact that hypnotic hallucinations are more vivid and that they compromise the sense of reality may explain why they involve greater top-down activation.

But why do hypnotic hallucinations engage with top-down mechanisms in a lateralised fashion? One possible explanation is that hypnotic hallucinations engage with top-down mechanisms in an incomplete or dissociated way. For example, it could be the case that while anterior areas activate posterior areas to induce and sustain the hallucinatory experience, they do not engage with feedback and control processes - hypnotic hallucinations may be experienced effortlessly and more vividly due to a lack of executive monitoring over early visual cortex. Myriad studies have shown a lateralised functioning of cognition in humans, with a strong correlation between executive function and activation in the left hemisphere in right-handed individuals (Barbey et al., 2012; Corballis, 2014; Fletcher et al., 2000; Gotts et al., 2013; Riès et al., 2016; Smith et al., 1996), including monitoring and feedback processing (Gruendler et al., 2011; Huster et al., 2009, 2011; Ocklenburg et al., 2011; Vallesi et al., 2009). This interpretation of our findings agrees with cold control theory, which proposes that hypnotic suggestions involve a disruption in executive monitoring and metacognition (Dienes et al., 2020; Dienes & Hutton, 2013; Dienes & Perner, 2007). Cold control theory, which stems from Higher-Order Thought theory of consciousness (Rosenthal, 2008, 2009), maintains that the experience of intention is mediated by the occurrence of higher-order thoughts, i.e. metacognitive states about one’s mental representations. These metacognitive states index authorship and other features of target first-order actions or representations. During hypnotic hallucination, these executive mechanisms of sense of agency and volition may be attenuated, thus explaining the experience of seeing a vivid face image that appeared by itself. In this light, hypnotic hallucination may be comparable to mental imagery in terms of their first-order representations whilst differing substantially in terms of their higher-order processes, including their conscious nature (Nanay, 2021).

Notably, many studies have found an association between visual hallucination and alterations in right-hemisphere function (Berthier & Starkstein, 1987; Jonas et al., 2014; Kim et al., 2019; Mégevand et al., 2014; Pakalnis et al., 1987; Sommer et al., 2008). Future studies should explore whether hypnotic hallucination engages with the same neural mechanisms of clinical hallucination. If so, hypnotic hallucination might be useful as a model to study psychotic hallucinations (Oakley & Halligan, 2013).

Our study presents several limitations. Firstly, even though we employed a large number of hypnotic virtuosos compared to past hypnosis studies, future studies should gather larger samples, based on power analyses. Secondly, we collected one vividness rating per participant in the hallucinatory condition. Future studies should collect subjective ratings on a trial-by-trial basis to capture statistical subtleties that our results may have missed. Additionally, future studies should explore other qualities of subjective experience during hypnotic hallucination. What makes a hallucinated image more vivid? Is it its visual clarity or level of detail? Or is it the experience of lack of control over their emergence? Future studies should delve into these phenomenological differences.

In conclusion, our findings suggest that hypnotic hallucination involves greater top-down processing through lateralised neural mechanisms than mental imagery, thus revealing important new insights into the neural mechanisms of hypnotic suggestion and hallucination.

## 7 Author contributions statement

R.C.L. and A.C-J., conceived the study; R.C.L., A.R-R., and A.C-J., developed the methodology; R.C.L., collected the experimental data; R.C.L., analysed the data; R.C.L., interpreted the findings; R.C.L., wrote the manuscript; A.R-R., D.H.,A.I., and A.C-J., constructively reviewed the manuscript.

## 8 Acknowledgements

The authors thank Johannes Fahrenfort for his technical advice; Tristan Bekinschtein, Verena Klar, Daniel Bor, and Valdas Noreika for suggestions on earlier versions of this manuscript; and Pedro Maldonado, José Luis Valdés, and Paul Delano for their suggestions at early stages of this project. R.C.L. thanks Verena Klar and Stephanie Schott for feeding him with amazing home-cooked meals during coronavirus lockdown Christmas 2020 at Oxford, UK – this article’s Introduction would not have been so efficiently drafted if he had relied on his own cooking skills instead.

