## Supplementary Material for "Hypnotic visual hallucination induces greater lateralised brain activity than visual imagery"

### SUPPLEMENTARY MATERIAL 1: Stimuli validation and standardisation

The validation and standardisation procedure of stimuli for imagery research in a Chilean population was based on Ganis & Schendan's (2008) adaptation paradigm and it is equivalent to the procedure carried out by Canales-Johnson et al. (2021).

#### Participants

A total of 60 undergraduate students, aged 18-36 years (*M*= 26.11 [*SD*= 3.21]), participated in the validation study – all from the cities of Santiago, Viña del Mar, and Valparaíso, Chile. They were asked to participate voluntarily either directly by the researchers or by public ads posted on their universities' bulletin boards.

#### Stimuli validation procedure

To validate facial images, we administered a questionnaire including the images of 125 celebrities (55 females, 70 males) to 60 university students who did not take part in the main study. They were asked to name each celebrity. Faces were considered valid when responses reached a minimum of 90% correct identification. Fifty-five percent of the pictures (i.e. 70) met this criterion and were thus selected to be used in the main study.

Given that the pictures were selected randomly, the recognition of the celebrities might have been affected by participants' age. To test this, we performed an ANOVA with age as a categorical factor, divided into the following age groups: 18-24 years (*M*=16.12 [5.16]); 25-30 years (*M*=16.53 [3.83]); and 31-36 years (*M*=16.75 [3.33]). No significant differences were found between these groups ($F_{(2.372)} = 0.71, p = .48$); suggesting that participants' age did not affect celebrity recognition.

We also selected 70 images of highly recognisable objects previously validated and standardised by Brodeur et al. (2010): the Bank of Standardized Stimuli (BOSS).

### Stimuli normalisation procedure

Seventy images of validated celebrity faces, and 70 images of objects were subsequently normalised based on size, brightness, and light intensity. They were converted to a greyscale image using Adobe® Photoshop®. To avoid potential distractions, only regions corresponding to eyes, nose, and mouth were selected for faces, keeping these facial areas inside an oval, as symmetrically as possible for each image (Supplementary Figure 1).

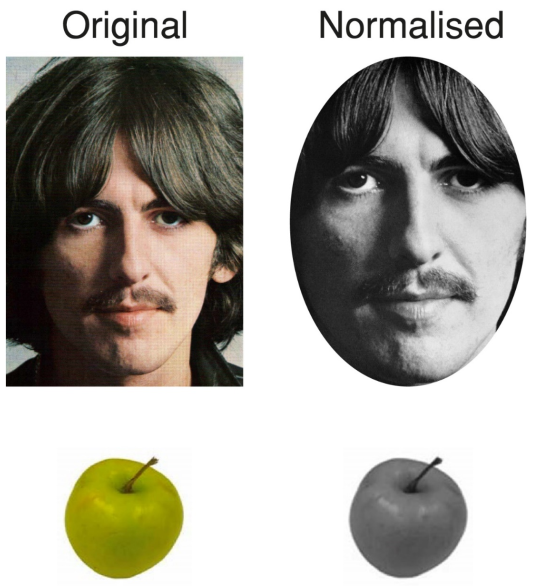


**Supplementary Figure 1.** Face and object photo normalisation. (Top) Typical celebrity photo used in the recognition test: the left image shows an example of those presented during the validation phase; the image on the right shows a normalised face ready to be used in the experimental task. (Bottom) Object photo selected from the BOSS and normalised as described above.

### Stimuli standardisation procedure

**Standardised face normalisation.** To determine whether the modifications described above affected image recognition, another recognition test was applied to a different set of 25 participants (*M*=23.1 [1.51]; 14 female). As before, faces that were recognised by 90% of participants were considered valid. All images met this criterion, and therefore no image was rejected. In addition, an interclass ANOVA was applied to examine the extent to which image variability affected face recognition variability. The result (0.05) indicated that only 5% of face recognition variability could be attributed to image variability after the normalisation process.

**Emotional valence recognition.** We tested whether the emotional valence of each face image was positive. Each face was classified within one of the following categories of emotional valence: positive (*M*=93.97 [7.59]), neutral (*M*=5.85 [7.49]) or negative (*M*=0.34 [1.64]). An ANOVA showed a main effect of category ($F_{(2.204)} = 4901.48, p < 0.001$). Post-hoc analysis (Tukey HSD test, MS=190.330,58, df=2.0) showed that the means of the faces with positive valence ($M$=93.62 [1.06]) were greater than faces with either neutral valence ($M$=88.11 [1.06]; $p < 0.01$) or with negative valence ($M$=93.62 [1.06]; $p < 0.01$). Recognition was considered valid since above 94% of the images were classified as having a positive emotional valence. Finally, we tested whether celebrity gender (male or female) affected the recognition of faces with positive valence. An ANOVA did not show a significant effect of gender ($F_{(1, 1.723)} = 0.41, p = 0.52$), suggesting that gender did not affect facial recognition of faces with positive valence.

**SUPPLEMENTARY MATERIAL 2: ERPs evoked by adaptor stimuli**

These analyses tested for differences in evoked ERPs to the adapted stimulus, i.e. time-locked to the keypress, not to test stimulus presentation. Therefore, they do not test our hypotheses but serve to ascertain that the N170/VPP shows its face-specific effect, i.e. more negative voltage to faces than to objects at both left and right N170 ROIs.

**P1 component**

The P1 component elicited by face and object stimuli peaked at 125 ms post-stimulus, in both imagery and perceptual conditions. P1 averaged voltage values were submitted to a 2 (stimulus category: face, object) × 2 (condition: imagery, perception) repeated-measures ANOVA. We found a main effect of condition ($F_{(1, 23)} = 22.745, p < .001, \eta p^2 = .497$), but not of stimulus ($F_{(1, 23)} = 0.669, p = .422, \eta p^2 = .028$), indicating that P1's voltage was significantly more positive in the perceptual condition ($M$ = 1.66 [1.5]) than in the imagery condition ($M$=0.498 [1.51]). The interaction did not reach significance ($F_{(1, 23)} = 2.182, p = .153, \eta p^2 = .087$). These results may reflect differences in visual sharpness and contrast between the imagined and perceived images.

**N170/VPP complex**

The N170/VPP complex elicited by face and object adaptor stimuli peaked around 181 ms post-keypress, in both imagery and perceptual conditions. The averaged voltage values were submitted to a 2 (stimulus category: face, object) × 2 (condition: imagery, perception) × 3 (ROI: left N170, right N170, VPP) repeated-measures ANOVA. We found a main effect of stimulus category ($F_{(1, 23)} = 9.104, p = .006, \eta p^2 = .284$), condition ($F_{(1, 23)} = 13.278, p < .001, \eta p^2 = .366$), and ROI ($F_{(1.255, 28.854)} = 6.057, p = .015, \eta p^2 = .208$). All the interactions reached significance: between stimulus category and condition ($F_{(1, 23)} = 8.735, p = .007, \eta p^2 = .275$), between stimulus category and ROI ($F_{(1.144, 26.306)} = 27.310, p < .001, \eta p^2 = .543$), between condition and ROI ($F_{(1.545, 35.533)} = 21.490, p < .001, \eta p^2 = .483$), and finally the three-way interaction ($F_{(1.388, 31.916)} = 23.434, p < .001, \eta p^2 = .505$). In summary, face stimuli evoked more negative voltage than object stimuli in the perceptual task but not in the imagery task on left and right N170 ROIs. The opposite was observed on VPP – more positive voltage evoked by face stimuli than object stimuli in the perceptual condition, thus we replicated this well-established finding (Bentin et al., 1996; Ganis & Schendan, 2008; Itier & Taylor, 2002; Joyce & Rossion, 2005; Kovács et al., 2006). These results suggest that the N170/VPP complex was evoked as expected in the perceptual condition, distinguishing between face and object images as expected. Results obtained from post hoc Bonferroni-corrected pairwise comparisons are summarised in tables, per interaction, as it follows: stimulus category and condition (Supplementary Table 1); stimulus category and ROI (Supplementary Table 2); condition and ROI (Supplementary Table 3); and the three-way interaction (Supplementary Table 4).

**Supplementary Table 1.**
Post hoc Bonferroni-corrected pairwise comparisons: Stimulus category * Condition interaction

| Factor 1 | Factor 2 | Mean Difference | SE | t | p |
|---|---|---|---|---|---|
| Face, Imagery | Object, Imagery | 0.029 | 0.123 | 0.237 | 1.000 |
| | Face, Perception | -0.447 | 0.219 | -2.046 | 0.293 |
| | Object, Perception | -0.966 | 0.214 | -4.516 | < .001 |
| Object, Imagery | Face, Perception | -0.477 | 0.214 | -2.228 | 0.201 |
| | Object, Perception | -0.995 | 0.219 | -4.553 | < .001 |
| Face, Perception | Object, Perception | -0.519 | 0.123 | -4.211 | < .001 |

**Supplementary Table 2.**
Post hoc Bonferroni-corrected pairwise comparisons: Stimulus category * Condition interaction

| Factor 1 | Factor 2 | Mean Difference | SE | t | p |
|---|---|---|---|---|---|
| Face, Left | Object, Left | -0.585 | 0.179 | -3.272 | 0.026 |
| | Face, Right | 0.241 | 0.420 | 0.574 | 1.000 |
| | Object, Right | -0.810 | 0.412 | -1.966 | 0.818 |
| | Face, Central | -1.934 | 0.420 | -4.609 | < .001 |
| | Object, Central | -1.032 | 0.412 | -2.504 | 0.231 |
| Object, Left | Face, Right | 0.826 | 0.412 | 2.003 | 0.753 |
| | Object, Right | -0.225 | 0.420 | -0.537 | 1.000 |
| | Face, Central | -1.349 | 0.412 | -3.272 | 0.028 |
| | Object, Central | -0.447 | 0.420 | -1.065 | 1.000 |
| Face, Right | Object, Right | -1.051 | 0.179 | -5.878 | < .001 |
| | Face, Central | -2.175 | 0.420 | -5.183 | < .001 |
| | Object, Central | -1.273 | 0.412 | -3.088 | 0.048 |
| Object, Right | Face, Central | -1.124 | 0.412 | -2.726 | 0.130 |
| | Object, Central | -0.222 | 0.420 | -0.529 | 1.000 |
| Face, Central | Object, Central | 0.902 | 0.179 | 5.044 | < .001 |

**Supplementary Table 3.**
Post hoc Bonferroni-corrected pairwise comparisons: Condition * ROI interaction

| Factor 1 | Factor 2 | Mean Difference | SE | t | p |
|---|---|---|---|---|---|
| Imagery, Left | Perception, Left | -1.760 | 0.357 | -4.927 | < .001 |
| | Imagery, Right | -0.057 | 0.473 | -0.120 | 1.000 |
| | Perception, Right | -1.688 | 0.467 | -3.612 | 0.008 |
| | Imagery, Central | -2.684 | 0.473 | -5.679 | < .001 |
| | Perception, Central | -1.457 | 0.467 | -3.119 | 0.038 |
| Perception, Left | Imagery, Right | 1.703 | 0.467 | 3.645 | 0.007 |
| | Perception, Right | 0.072 | 0.473 | 0.153 | 1.000 |
| | Imagery, Central | -0.924 | 0.467 | -1.978 | 0.773 |
| | Perception, Central | 0.303 | 0.473 | 0.640 | 1.000 |
| Imagery, Right | Perception, Right | -1.631 | 0.357 | -4.565 | < .001 |
| | Imagery, Central | -2.627 | 0.473 | -5.559 | < .001 |
| | Perception, Central | -1.401 | 0.467 | -2.997 | 0.055 |
| Perception, Right | Imagery, Central | -0.996 | 0.467 | -2.132 | 0.542 |
| | Perception, Central | 0.230 | 0.473 | 0.487 | 1.000 |
| Imagery, Central | Perception, Central | 1.227 | 0.357 | 3.434 | 0.015 |

**Supplementary Table 4.**
Post hoc Bonferroni-corrected pairwise comparisons: Stimulus category * Condition * ROI interaction

| Factor 1 | Factor 2 | Mean Difference | SE | t | p |
|---|---|---|---|---|---|
| Face, Imagery, Left | Object, Imagery, Left | 0.163 | 0.270 | 0.604 | 1.000 |
| | Face, Perception, Left | -1.011 | 0.411 | -2.463 | 1.000 |
| | Object, Perception, Left | -2.345 | 0.399 | -5.870 | < .001 |
| | Face, Imagery, Right | 0.039 | 0.517 | 0.075 | 1.000 |
| | Object, Imagery, Right | 0.011 | 0.503 | 0.021 | 1.000 |
| | Face, Perception, Right | -0.569 | 0.504 | -1.128 | 1.000 |
| | Object, Perception, Right | -2.643 | 0.506 | -5.228 | < .001 |
| | Face, Imagery, Central | -2.579 | 0.517 | -4.992 | < .001 |
| | Object, Imagery, Central | -2.626 | 0.503 | -5.221 | < .001 |
| | Face, Perception, Central | -2.302 | 0.504 | -4.566 | < .001 |
| | Object, Perception, Central | -0.450 | 0.506 | -0.890 | 1.000 |
| Object, Imagery, Left | Face, Perception, Left | -1.175 | 0.399 | -2.941 | 0.268 |
| | Object, Perception, Left | -2.509 | 0.411 | -6.108 | < .001 |
| | Face, Imagery, Right | -0.124 | 0.503 | -0.248 | 1.000 |
| | Object, Imagery, Right | -0.153 | 0.517 | -0.295 | 1.000 |
| | Face, Perception, Right | -0.732 | 0.506 | -1.448 | 1.000 |
| | Object, Perception, Right | -2.806 | 0.504 | -5.567 | < .001 |
| | Face, Imagery, Central | -2.742 | 0.503 | -5.451 | < .001 |
| | Object, Imagery, Central | -2.789 | 0.517 | -5.401 | < .001 |
| | Face, Perception, Central | -2.465 | 0.506 | -4.875 | < .001 |
| | Object, Perception, Central | -0.613 | 0.504 | -1.216 | 1.000 |
| Face, Perception, Left | Object, Perception, Left | -1.334 | 0.270 | -4.934 | < .001 |
| | Face, Imagery, Right | 1.050 | 0.504 | 2.083 | 1.000 |
| | Object, Imagery, Right | 1.022 | 0.506 | 2.022 | 1.000 |

**Supplementary Table 4.**
Post hoc Bonferroni-corrected pairwise comparisons: Stimulus category * Condition * ROI interaction

| Factor 1 | Factor 2 | Mean Difference | SE | t | p |
|---|---|---|---|---|---|
| | Face, Perception, Right | 0.443 | 0.517 | 0.857 | 1.000 |
| | Object, Perception, Right | -1.632 | 0.503 | -3.244 | 0.106 |
| | Face, Imagery, Central | -1.567 | 0.504 | -3.109 | 0.160 |
| | Object, Imagery, Central | -1.615 | 0.506 | -3.194 | 0.123 |
| | Face, Perception, Central | -1.290 | 0.517 | -2.498 | 0.924 |
| | Object, Perception, Central | 0.562 | 0.503 | 1.116 | 1.000 |
| Object, Perception, Left | Face, Imagery, Right | 2.384 | 0.506 | 4.715 | < .001 |
| | Object, Imagery, Right | 2.356 | 0.504 | 4.673 | < .001 |
| | Face, Perception, Right | 1.776 | 0.503 | 3.531 | 0.041 |
| | Object, Perception, Right | -0.298 | 0.517 | -0.577 | 1.000 |
| | Face, Imagery, Central | -0.233 | 0.506 | -0.462 | 1.000 |
| | Object, Imagery, Central | -0.281 | 0.504 | -0.557 | 1.000 |
| | Face, Perception, Central | 0.044 | 0.503 | 0.087 | 1.000 |
| | Object, Perception, Central | 1.895 | 0.517 | 3.669 | 0.025 |
| Face, Imagery, Right | Object, Imagery, Right | -0.028 | 0.270 | -0.104 | 1.000 |
| | Face, Perception, Right | -0.608 | 0.411 | -1.479 | 1.000 |
| | Object, Perception, Right | -2.682 | 0.399 | -6.713 | < .001 |
| | Face, Imagery, Central | -2.617 | 0.517 | -5.067 | < .001 |
| | Object, Imagery, Central | -2.665 | 0.503 | -5.298 | < .001 |
| | Face, Perception, Central | -2.340 | 0.504 | -4.643 | < .001 |
| | Object, Perception, Central | -0.489 | 0.506 | -0.967 | 1.000 |
| Object, Imagery, Right | Face, Perception, Right | -0.580 | 0.399 | -1.451 | 1.000 |
| | Object, Perception, Right | -2.654 | 0.411 | -6.462 | < .001 |
| | Face, Imagery, Central | -2.589 | 0.503 | -5.148 | < .001 |
| | Object, Imagery, Central | -2.637 | 0.517 | -5.105 | < .001 |
| | Face, Perception, Central | -2.312 | 0.506 | -4.573 | < .001 |
| | Object, Perception, Central | -0.461 | 0.504 | -0.914 | 1.000 |
| Face, Perception, Right | Object, Perception, Right | -2.074 | 0.270 | -7.674 | < .001 |
| | Face, Imagery, Central | -2.010 | 0.504 | -3.987 | 0.008 |
| | Object, Imagery, Central | -2.057 | 0.506 | -4.069 | 0.006 |
| | Face, Perception, Central | -1.733 | 0.517 | -3.355 | 0.072 |
| | Object, Perception, Central | 0.119 | 0.503 | 0.236 | 1.000 |
| Object, Perception, Right | Face, Imagery, Central | 0.065 | 0.506 | 0.128 | 1.000 |
| | Object, Imagery, Central | 0.017 | 0.504 | 0.034 | 1.000 |
| | Face, Perception, Central | 0.342 | 0.503 | 0.679 | 1.000 |
| | Object, Perception, Central | 2.193 | 0.517 | 4.246 | 0.003 |
| Face, Imagery, Central | Object, Imagery, Central | -0.048 | 0.270 | -0.176 | 1.000 |
| | Face, Perception, Central | 0.277 | 0.411 | 0.675 | 1.000 |
| | Object, Perception, Central | 2.129 | 0.399 | 5.328 | < .001 |
| Object, Imagery, Central | Face, Perception, Central | 0.325 | 0.399 | 0.813 | 1.000 |
| | Object, Perception, Central | 2.176 | 0.411 | 5.299 | < .001 |
| Face, Perception, Central | Object, Perception, Central | 1.852 | 0.270 | 6.850 | < .001 |

### SUPPLEMENTARY MATERIAL 3: N170/VPP evoked by test stimuli in hallucinators

Does hypnotic hallucination of faces modulate the N170/VPP complex differently than visual imagery and perception of faces? We submitted the data to a 3 (condition: hallucination, imagery, perception) × 3 (ROI: left N170, right N170, VPP) repeated-measures ANOVA (Figure 5). We found a main effect of condition ($F_{(1.56,\ 9.358)} = 10.532, p = .006, \eta p^2 = .637$) and ROI ($F_{(1.904,\ 11.425)} = 34.314, p < .001, \eta p^2 = .851$). The interaction also reached significance ($F_{(2.299,\ 13.794)} = 31.39, p < .001, \eta p^2 = .84$). Bonferroni-corrected pairwise comparisons revealed that voltage values in the hallucinatory condition (*M*=-2.265 [1.324]) were more negative than in the perceptual condition at left N170 (*M*=-0.132 [0.705]), ($t = -4.949, p < .001$). Likewise, voltage values in the imagery condition (*M*=-1.819 [1.039]) were more negative than in the perceptual condition at the same ROI ($t = -3.915, p = .014$). However, the hallucinatory condition and the imagery condition did not differ significantly at left N170 ($t = -1.034, p > .1$). Crucially, however, we found that voltage values in the hallucinatory condition (*M*=-4.166 [0.742]) were significantly more negative than in the imagery condition (*M*=-2.623 [1.136]), ($t = -3.58, p = .036$) at right N170, and that the voltage values in the latter were significantly more negative than in the perceptual condition (*M*=-0.415 [1.045]), ($t = -5.125, p < .001$). Therefore, we found that hypnotic hallucination differs from mental imagery in how it modulates the N170/VPP complex only at the right hemisphere's N170 component – this is our main finding. Supplementary Table 5 contains the results obtained from post hoc Bonferroni-corrected pairwise comparisons followed by the interaction between Condition and ROI.

**Supplementary Table 5.**
Post hoc Bonferroni-corrected pairwise comparisons: Condition * ROI interaction

| | | Mean Difference | SE | t | p |
|---|---|---|---|---|---|
| Hallucination, Left | Imagery, Left | -0.446 | 0.431 | -1.034 | 1.000 |
| | Perception, Left | -2.133 | 0.431 | -4.949 | < .001 |
| | Hallucination, Right | 1.902 | 0.632 | 3.007 | 0.221 |
| | Imagery, Right | 0.359 | 0.623 | 0.575 | 1.000 |
| | Perception, Right | -1.850 | 0.623 | -2.968 | 0.245 |
| | Hallucination, Central | -4.940 | 0.632 | -7.812 | < .001 |
| | Imagery, Central | -4.732 | 0.623 | -7.591 | < .001 |
| | Perception, Central | -2.261 | 0.623 | -3.626 | 0.050 |
| Imagery, Left | Perception, Left | -1.687 | 0.431 | -3.915 | 0.014 |
| | Hallucination, Right | 2.347 | 0.623 | 3.765 | 0.035 |
| | Imagery, Right | 0.804 | 0.632 | 1.272 | 1.000 |
| | Perception, Right | -1.405 | 0.623 | -2.253 | 1.000 |
| | Hallucination, Central | -4.495 | 0.623 | -7.210 | < .001 |
| | Imagery, Central | -4.286 | 0.632 | -6.779 | < .001 |
| | Perception, Central | -1.815 | 0.623 | -2.912 | 0.279 |
| Perception, Left | Hallucination, Right | 4.035 | 0.623 | 6.472 | < .001 |
| | Imagery, Right | 2.491 | 0.623 | 3.997 | 0.020 |
| | Perception, Right | 0.283 | 0.632 | 0.447 | 1.000 |
| | Hallucination, Central | -2.807 | 0.623 | -4.503 | 0.006 |
| | Imagery, Central | -2.599 | 0.623 | -4.169 | 0.013 |

**Supplementary Table 5.**
Post hoc Bonferroni-corrected pairwise comparisons: Condition * ROI interaction

| | | Mean Difference | SE | t | p |
|---|---|---|---|---|---|
| | Perception, Central | -0.128 | 0.632 | -0.202 | 1.000 |
| Hallucination, Right | Imagery, Right | -1.543 | 0.431 | -3.580 | 0.036 |
| | Perception, Right | -3.752 | 0.431 | -8.705 | < .001 |
| | Hallucination, Central | -6.842 | 0.632 | -10.820 | < .001 |
| | Imagery, Central | -6.634 | 0.623 | -10.641 | < .001 |
| | Perception, Central | -4.162 | 0.623 | -6.677 | < .001 |
| Imagery, Right | Perception, Right | -2.209 | 0.431 | -5.125 | < .001 |
| | Hallucination, Central | -5.299 | 0.623 | -8.500 | < .001 |
| | Imagery, Central | -5.091 | 0.632 | -8.050 | < .001 |
| | Perception, Central | -2.619 | 0.623 | -4.202 | 0.012 |
| Perception, Right | Hallucination, Central | -3.090 | 0.623 | -4.957 | 0.002 |
| | Imagery, Central | -2.882 | 0.623 | -4.623 | 0.004 |
| | Perception, Central | -0.411 | 0.632 | -0.649 | 1.000 |
| Hallucination, Central | Imagery, Central | 0.208 | 0.431 | 0.483 | 1.000 |
| | Perception, Central | 2.679 | 0.431 | 6.217 | < .001 |
| Imagery, Central | Perception, Central | 2.471 | 0.431 | 5.734 | < .001 |

### SUPPLEMENTARY MATERIAL 4: N170/VPP evoked by test stimuli in non-hallucinators

Do hypnotic virtuosos who do not hallucinate show this lateralised effect of hallucination on N170? We submitted the non-hallucinator participants' data to a 3 (condition: hallucination, imagery, perception) × 3 (ROI: left N170, right N170, VPP) repeated-measures ANOVA (Figure 6). We found a main effect of condition ($F_{(1.323,\ 7.941)} = 5.224, p = .045, \eta p^2 = .465$) and ROI ($F_{(1.179,\ 7.075)} = 8.388, p = .02, \eta p^2 = .583$). The interaction also reached significance ($F_{(2.413,\ 14.48)} = 14.532, p < .001, \eta p^2 = .708$). Bonferroni-corrected pairwise comparisons revealed that voltage values in the hallucinatory condition (*M*=-2.336 [1.322]) were more negative than in the perceptual condition at left N170 (*M*=-0.358 [1.305]), ($t = -4.171, p = .007$). Likewise, voltage values in the imagery condition (*M*=-2.277 [1.566]) were more negative than in the perceptual condition at the same ROI ($t = -4.047, p = .001$). However, the hallucinatory condition and the imagery condition did not differ significantly at left N170 ($t = -0.124, p > .1$). We found similar results at right N170. Voltage values in the hallucinatory condition (*M*=-2.588 [2.587]) were more negative than in the perceptual condition (*M*=-0.753 [1.852]), ($t = -3.871, p = .017$) Similarly, voltage values in the imagery condition (*M*=-2.988 [1.964]) were more negative than in the perceptual condition at right N170 ($t = -4.714, p = .001$). Notably, however, we did not find a significant difference between hallucinatory and imagery conditions at right N170 ($t = 0.843, p > .1$). Therefore, unlike the hallucinator group, who did exhibit such a difference at right N170, non-hallucinators did not. Finally, we estimated Bayes factors to assess this null effect between hallucinatory and imagery conditions at right N170, which provided anecdotal evidence in favour of the null hypothesis ($BF_{01} = 2.274$).

Altogether, these results may suggest that the ability to hallucinate faces as a response to posthypnotic suggestions is associated with a lateralised effect at the N170 component – while hallucinators exhibit a modulation at right N170 between hallucinatory and imagery conditions, non-hallucinators do not exhibit this modulation. Supplementary Table 6 contains the results obtained from post hoc Bonferroni-corrected pairwise comparisons followed by the interaction between Condition and ROI.

**Supplementary Table 6.**

Post hoc Bonferroni-corrected pairwise comparisons: Condition * ROI interaction

| Factor 1 | Factor 2 | Mean Difference | SE | t | p |
|---|---|---|---|---|---|
| Hallucination, Left | Imagery, Left | -0.059 | 0.474 | -0.124 | 1.000 |
| | Perception, Left | -1.978 | 0.474 | -4.171 | 0.007 |
| | Hallucination, Right | 0.253 | 1.199 | 0.211 | 1.000 |
| | Imagery, Right | 0.652 | 1.184 | 0.551 | 1.000 |
| | Perception, Right | -1.583 | 1.184 | -1.337 | 1.000 |
| | Hallucination, Central | -4.888 | 1.199 | -4.078 | 0.034 |
| | Imagery, Central | -5.340 | 1.184 | -4.511 | 0.016 |
| | Perception, Central | -3.032 | 1.184 | -2.561 | 0.790 |
| Imagery, Left | Perception, Left | -1.919 | 0.474 | -4.047 | 0.010 |
| | Hallucination, Right | 0.312 | 1.184 | 0.263 | 1.000 |
| | Imagery, Right | 0.711 | 1.199 | 0.593 | 1.000 |
| | Perception, Right | -1.524 | 1.184 | -1.288 | 1.000 |
| | Hallucination, Central | -4.829 | 1.184 | -4.079 | 0.037 |
| | Imagery, Central | -5.281 | 1.199 | -4.406 | 0.017 |
| | Perception, Central | -2.974 | 1.184 | -2.512 | 0.872 |
| Perception, Left | Hallucination, Right | 2.231 | 1.184 | 1.884 | 1.000 |
| | Imagery, Right | 2.630 | 1.184 | 2.222 | 1.000 |
| | Perception, Right | 0.395 | 1.199 | 0.329 | 1.000 |
| | Hallucination, Central | -2.910 | 1.184 | -2.458 | 0.968 |
| | Imagery, Central | -3.362 | 1.184 | -2.840 | 0.454 |
| | Perception, Central | -1.054 | 1.199 | -0.880 | 1.000 |
| Hallucination, Right | Imagery, Right | 0.400 | 0.474 | 0.843 | 1.000 |
| | Perception, Right | -1.836 | 0.474 | -3.871 | 0.017 |
| | Hallucination, Central | -5.141 | 1.199 | -4.289 | 0.022 |
| | Imagery, Central | -5.593 | 1.184 | -4.724 | 0.010 |
| | Perception, Central | -3.285 | 1.184 | -2.775 | 0.517 |
| Imagery, Right | Perception, Right | -2.236 | 0.474 | -4.714 | 0.001 |
| | Hallucination, Central | -5.540 | 1.184 | -4.680 | 0.011 |
| | Imagery, Central | -5.993 | 1.199 | -4.999 | 0.005 |
| | Perception, Central | -3.685 | 1.184 | -3.112 | 0.262 |
| Perception, Right | Hallucination, Central | -3.305 | 1.184 | -2.792 | 0.500 |
| | Imagery, Central | -3.757 | 1.184 | -3.173 | 0.231 |
| | Perception, Central | -1.449 | 1.199 | -1.209 | 1.000 |
| Hallucination, Central | Imagery, Central | -0.452 | 0.474 | -0.953 | 1.000 |
| | Perception, Central | 1.856 | 0.474 | 3.913 | 0.015 |
| Imagery, Central | Perception, Central | 2.308 | 0.474 | 4.866 | < .001 |
